# Early life thermal plasticity and adaptive divergence among populations of Arctic charr (*Salvelinus alpinus*)

**DOI:** 10.64898/2026.03.06.710097

**Authors:** Hervé Rogissart, Lisandrina Mari, Guillaume Evanno, Martin Daufresne, Luca Fumagalli, Jean Guillard, Allan Raffard, Emilien Lasne

**Affiliations:** Univ. Savoie Mont Blanc, INRAE, CARRTEL, Thonon-les-Bains, France; Department of Biological and Environmental Science, University of Jyväskylä, Jyväskylä, Finland; DECOD (Ecosystem Dynamics and Sustainability), INRAE, Institut Agro, IFREMER, Rennes, France; INRAE, Aix-Marseille Univ., RECOVER, Aix-en-Provence, France; Laboratory for Conservation Biology, Department of Ecology and Evolution, Biophore, University of Lausanne, Lausanne, Switzerland; Swiss Human Institute of Forensic Taphonomy, University Centre of Legal Medicine Lausanne-Geneva, Lausanne University Hospital and University of Lausanne, Switzerland

**Keywords:** phenotypic plasticity, quantitative genetics, adaptive divergence, reaction norm, common garden

## Abstract

Climate warming alters the thermal environment experienced by ectotherms, whose physiological performance and fitness are constrained by temperature. Early life stages are often the temperature-sensitive phases of the life cycle, particularly in freshwater stenotherms such as the Arctic charr (*Salvelinus alpinus*). In this study, we used a common garden approach to compare the response and phenotypic plasticity of four charr populations to warmer conditions. These populations inhabit thermally contrasted lakes, including both native and introduced populations. We reared embryos at either an optimal (5°C) or a warmer (8.5°C) temperature. We tested adaptive divergence among populations in four traits (survival, incubation duration, body length and yolk sac volume) using *Q_st_* − *F_st_* comparisons. We found that warmer temperature reduced body size, survival and incubation time, while thermal reaction norms differed among populations. Unexpectedly, populations originating from warmer environments did not consistently exhibit higher trait values under elevated temperatures. In contrast, the colder high-altitude population exhibited higher survival rates and lower yolk reserves for a given size under heat stress than the other populations. Our results highlight substantial variation among populations in responses to heat stress, emphasizing the complexity of predicting population responses to climate warming.

## Introduction

Population persistence in warming environments depends on phenotypic change driven by both genetic evolution and phenotypic plasticity (Chevin & Bridle, 2025). Phenotypic plasticity, the ability of a genotype to produce various phenotypes under different environmental conditions, is particularly important during early development, since thermal sensitivity shapes growth and survival trajectories with potential carry-over effects on later life-history traits (Pankhurst & Munday, 2011; Massey & Hutchings, 2021; Jonsson *et al*., 2022). Genetic adaptation can occur at longer timescales when traits evolve towards distinct optima through time or in different habitats (Stearns, 1992; Gienapp *et al*., 2008). Phenotypic plasticity can also evolve when there is genetic variation in plastic responses, resulting in significant genotype-by-environment (*G* × *E*) interactions (Ghalambor *et al*., 2007; Saltz *et al*., 2018). Such interactions can reveal population-specific reaction norms consistent with local adaptation to contrasting environments (Hutchings, 2011; Oomen & Hutchings, 2015). Understanding the mechanisms that drive divergence among populations is crucial to predict how vulnerable, isolated populations may respond to future environmental changes (Kawecki & Ebert, 2004; Hendry *et al*., 2011; Urban *et al*., 2024). Adaptive divergence can be assessed by comparing quantitative trait differentiation (*Q_st_*) with neutral genetic differentiation (*F_st_*) (Merilä & Crnokrak, 2001; Leinonen *et al*., 2013). Briefly, this comparison provides a means to disentangle the causes of population divergence, with *Q_st_* values higher, lower, or similar to *F_st_* indicating divergent selection, stabilizing selection, or genetic drift, respectively (Leinonen *et al*., 2013).

In the context of climate change, ectotherms represent particularly relevant models for studying the mechanisms underlying microevolution and adaptation to rapid environmental changes (Angilletta *et al*., 2002; Narum *et al*., 2013). Most aquatic organisms are obligate poikilotherms, meaning they must cope with the thermal environment that influences their body temperature and all subsequent biological processes (Coutant, 1976; Pörtner & Farrell, 2008). In fishes, temperature affects a wide range of fitness-related traits, including embryonic survival, developmental rates, growth, metabolism, energy allocation (Larsson & Berglund, 2005; Hemmer-Brepson *et al*., 2013), and reproductive capacity (King *et al*., 2007; Pankhurst & King, 2010). Salmonids exhibit strong physiological sensitivity to temperature variation, especially during embryonic and larval development, making them well suited to investigate thermal adaptation and plasticity (Elliott & Elliott, 2010; Kavanagh *et al*., 2010; Jonsson, 2023; Gendron *et al*., 2025).

Early life stages of salmonids (embryonic and alevin), in particular, represent critical periods of development and are often the most sensitive to temperature change (Jeuthe *et al*., 2016; O’Dea *et al*., 2019; Dahlke *et al*., 2020, 2022). During embryogenesis and early larval development, organisms undergo rapid cellular differentiation and organogenesis while possessing a limited capacity for behavioral thermoregulation, making them strongly dependent on local thermal conditions (Blaxter, 1991). Temperature experienced during these stages can negatively influence survival, developmental rate, larval and emergence size stage (Realis-Doyelle *et al*., 2018; Mari *et al*., 2021). Consequently, small changes in developmental temperature can induce marked effects on early-life traits and survival from egg to larval stages, with potential long-term consequences for population persistence (Elliott & Elliott, 2010; Dahlke *et al*., 2020; Takatsu *et al*., 2022).

Many salmonid populations are currently experiencing rapid declines, particularly in cold-adapted freshwater systems, raising concerns about the limits of adaptive and plastic responses under ongoing climate change (Muhlfeld *et al*., 2019; Barley *et al*., 2021; Williams *et al*., 2024). Due to its climate and topography, the Alpine region is severely exposed to climate change-induced temperature increases, and warming is projected to intensify throughout the 21^st^ century (Bogataj, 2007; Dumont *et al*., 2025), with implications for cold-adapted species such as Arctic charr, *Salvelinus alpinus* (Linnaeus, 1758). In Arctic charr lakes, temperatures on spawning grounds can exceed 8 °C during embryonic development (Mari *et al*., 2021). Such temperatures are known to affect early developmental performance (Swift, 1965; Elliott & Elliott, 2010). Given the occurrence of higher temperatures in peri-Alpine lakes compared to Arctic charr habitats in colder environments at higher latitudes or altitudes, local adaptation to temperature at low elevation at the southern limit of its range is likely. Knowledge on the evolutionary potential of species inhabiting peri-Alpine lakes (large lakes located along the margins of the Alps), and mountain lakes (> 1000 m a.s.l.) located in the Alps and French Massif Central is critical for evaluating the implications of global change in this region.

Here, we investigate the plasticity and divergence among Arctic charr populations from Alpine and peri-Alpine lakes. In Europe, the peri-Alpine lakes region constitutes the natural southern edge of its range, and many populations were introduced into high-elevation Alpine lakes over the past two centuries (Machino, 1991). Because population history can influence patterns of quantitative trait divergence and genotype-by-environment interactions, native and introduced populations may differ in their early-life responses to thermal conditions (Koskinen *et al*., 2002; Kavanagh *et al*., 2010; Christie *et al*., 2012, 2014; Dayan *et al*., 2024). Previous studies indicate very narrow thermal tolerance ranges for the early life stages of the species, with reported optimal egg incubation temperatures generally between 6 and 8°C (Elliott & Elliott, 2010; Sæther *et al*., 2016). However, most investigations have been conducted on hatchery stocks (Swift, 1965; Jungwirth & Winkler, 1984; De March, 1995; Bebak *et al*., 2000; Jeuthe *et al*., 2016; Olk *et al*., 2020), which may not fully reflect the responses of wild populations. Information on wild populations remains scarce.

The goal of this study was to test whether early life-history traits exhibit adaptive divergence and population-specific responses to thermal stress among four wild Arctic charr populations. We selected two native populations from Lakes Geneva and Constance and two introduced populations from Lakes Allos and Pavin. These populations form two native– introduced pairs (Geneva–Allos and Constance–Pavin), providing a framework to examine divergence in early-life traits under contrasting environmental conditions. Lakes Geneva and Constance are large peri-Alpine lakes subjected to long histories of human exploitation, whereas Lakes Allos and Pavin are small mountain lakes characterized by colder and more oligotrophic conditions with minimal anthropogenic pressure. To disentangle genetic and environmental components of early-life trait variation, we conducted a common-garden experiment in which larvae from all populations were reared under contrasting thermal conditions (Kawecki & Ebert, 2004). Specifically, we investigated phenotypic plasticity and *G* × *E* interactions in thermal reaction norms across two native-introduced population pairs. We then compared neutral genetic differentiation among populations using microsatellite markers and used *Q_st_* − *F_st_* an approach to test whether divergence in early life-history traits exceeded neutral expectations, thereby providing insight into the relative contributions of adaptive and neutral processes.

## Materials and Methods

### Model species

The Arctic charr is a cold-water salmonid, primarily freshwater but facultatively anadromous, with a boreal and circumpolar distribution (Klemetsen, 2013; Weinstein *et al*., 2024). As the world’s northernmost freshwater fish species, Arctic charr is considered particularly vulnerable to climate warming (Lehtonen, 1998; Muhlfeld *et al*., 2024). Elevated temperatures can affect growth, survival, reproduction and early developmental processes (Jeuthe *et al*., 2016; Kelly *et al*., 2020; Gendron *et al*., 2025). The species reaches its southernmost native range limit in deep peri-Alpine lakes of the Alpine arc (Machino, 1991; Alexander & Seehausen, 2021). In such lakes, reproduction generally occurs between November and January, under cold-water conditions typically ranging from 4°C to 8°C (Guillard *et al*., 1992). Spawning ground water temperatures vary significantly among lakes. In Lake Geneva, spawning temperatures usually range from 7-8°C and may occasionally exceed 8 °C (Mari *et al*., 2016), whereas in Lake Constance they range from 5-6°C (Figure 1). In the high-elevation lakes such as Allos and Pavin, mean winter water temperatures between December and March average 3-4°C (Figure 1).

**Figure 1.**
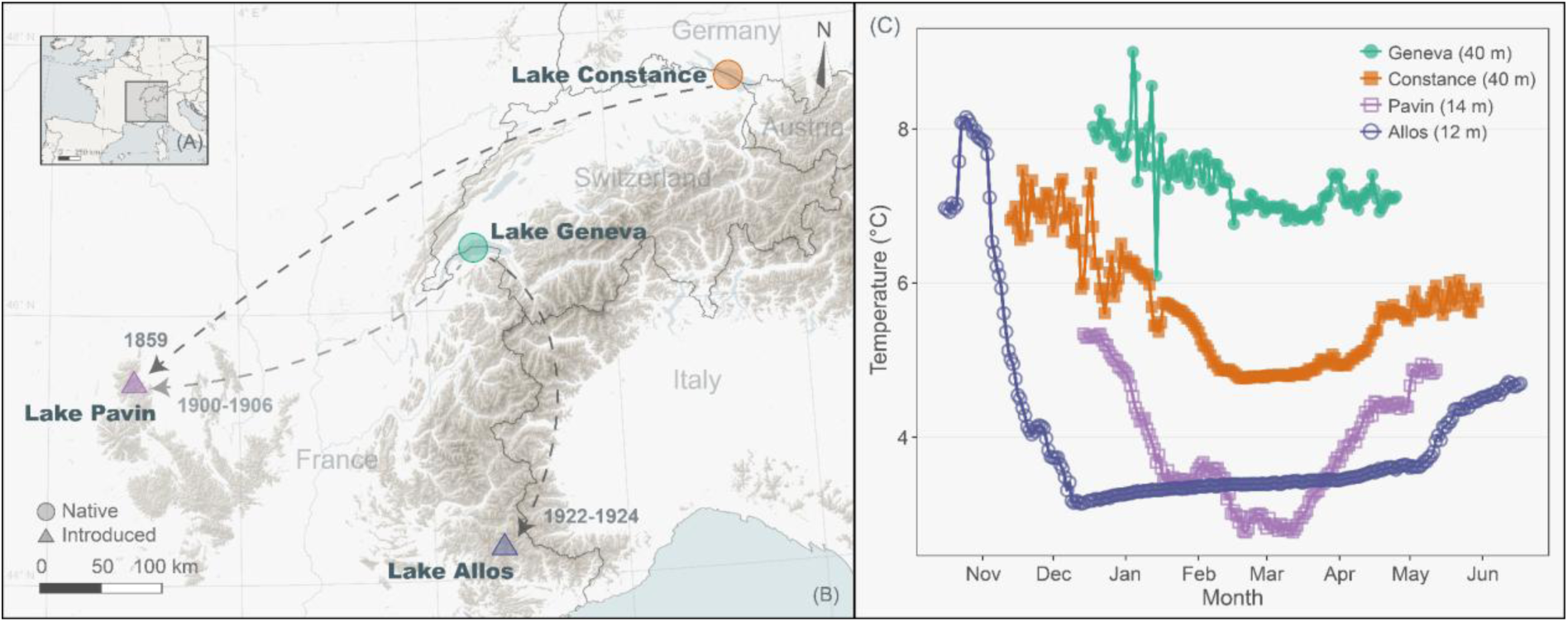
A, location of the study area within Western Europe. B, location of the four-study site, Lake Constance (Germany/Switzerland), Lake Geneva (France/Switzerland), Lake Pavin (France) and Lake Allos (France). White dashed arrows indicate the documented introduction of Arctic charr (*Salvelinus alpinus*) from Lake Constance to Pavin in 1859, from Lake Geneva to Lake Pavin in 1900 and 1906, and from Lake Geneva to Lake Allos in 1922 and 1924 (Machino & Rivier, 2002). Circles indicate native populations (Geneva and Constance), and triangles indicate introduced populations (Allos and Pavin). Digital elevation model data from (de Ferranti, 2022). C, winter temperature profiles at depths corresponding to Arctic charr spawning grounds in each lake, based on observations collected between 2014 and 2016.

### Study populations and historical background

This study focuses on four wild Arctic charr populations located in contrasting lakes that span a pronounced altitudinal and thermal gradient, providing naturally different environmental conditions (Figure 1, Table S1). The Arctic charr is native to lakes Geneva and Constance, where the populations became landlocked following the retreat of glaciers approximately 13,000 years ago (Dussart, 1955). These populations are still stocked annually through supportive breeding, using juveniles produced in hatcheries from local wild-caught spawners originating from the same population and released into the wild at the first-feeding stage (Caudron *et al*., 2014). Although occasional allochthonous stockings have occurred in the past, previous research indicated that these introductions had limited effects on the genetic diversity (Brunner *et al*., 2001) and did not completely alter the genetic integrity of these populations (Savary *et al*., 2017).

In contrast, Arctic charr were introduced to Lakes Pavin and Allos during the late 19^th^ and early 20^th^ centuries. In Lake Pavin, introductions occurred in 1859 using 8,000 alevins from the Huningue hatchery (Alsace, France) (Machino, 1991; Desmolles, 2016). This hatchery originally reared fish from wild stocks of the Rhine basin (Switzerland, Wurtemberg, and Bavaria; Coumes, 1862), followed by a second introduction between 1900 and 1906 using 2,500 eyed eggs from Lake Geneva (Crettiez, 1906; Reynouard, 1909; Machino, 1991; Desmolles, 2016). The Pavin population has since been managed by supportive breeding, with no further allochthonous introduction documented. In Lake Allos, Arctic charr were first introduced in 1922 using 2,500 alevins originating from Lake Geneva, followed by a second and last introduction in 1924 (Machino & Rivier, 2002).

### Sampling design and rearing conditions

Sampling and fertilizations were conducted during the winter of 2016, between mid-November and early December, corresponding to the natural spawning season (Table S1). Adults were captured on spawning grounds using benthic gill nets. For each population, individuals were crossed using block mating designs with two to three females and three to four males per block (Table S2), depending on the availability of mature individuals and female fecundity. In total, 124 families were established (Table S2). All spawners were anesthetized with 0.35 mL eugenol.L^–1^ (i.e., clove oil diluted 1:10 in 95% ethanol), measured, weighed, and pectoral fin clips were collected for genotyping. Thirty ova per female were collected and dry-weighed to estimate mean egg size per family. All adults were handled briefly and released.

Ova from the Geneva, Constance, and Pavin populations were fertilized in hatchery facilities located near each lake. During fertilization and transport to the INRAE facilities in Thonon-les-Bains, water temperature was kept between 2 and 5 °C. For logistical reasons, Allos spawners were transported to the INRAE facilities, where fertilization was performed. Following fertilization, eggs (n=8,628) were reared singly into 24-well cell culture plates (Nunc MicroWell^TM^, Thermo Fisher Scientific), targeting three replicates of 12 eggs (N = 36 individuals per family per treatment). The planned allocation was achieved for most families (111/124, 90%), whereas the remaining 13 families had slightly lower total egg numbers (39-70 eggs per family), reflecting variation in female fecundity, while remaining evenly distributed between temperature treatments. Eggs were incubated in thermal chambers at either 5 °C (4.996 ± 0.049 °C), referred to as cold treatment, or 8.5 °C (8.493 ± 0.038 °C), referred to as warm treatment. Temperature was recorded every 30-min using thermal sensors (HOBO UX100-001, Onset Computer Co., USA) placed in both chambers. Unfertilized eggs and early embryo mortality were recorded at the eyed stage.

Embryos were then monitored daily for mortality and hatching. Each well contained 2 mL of mineral water, and plates were covered with lids allowing gas exchange. Water was not renewed during incubation (water level remained constant throughout the incubation). Because hatched larvae could not be maintained in 24-well plates until yolk sac resorption, all live hatchlings from the same family were pooled after hatching, split into two replicates, and transferred to 50 x 50 mm incubators within a recirculating water system. Families were randomized within the system and exposed to identical water conditions. This transfer occurred five to nine days after the first hatching event was recorded within each population. Larvae were subsequently monitored daily for mortality, and dead individuals were removed. The experiment ended near yolk sac resorption, approximately 148 days post-fertilization (195–205 accumulated degree days, ADD calculated as the cumulative sum of daily mean water temperatures during the time elapsed) at cold and 100 days post-fertilization (215–230 ADD) at warm.

### Early-life history traits

Survival at hatching was estimated as a binary response (0 = did not hatch, 1 = successfully hatched) among embryos that reached the eyed stage. Incubation duration was expressed in ADD. Body length measurements were taken at two timepoints: *(i)* T1 at hatching immediately after transfer to the recirculating water system (aged between 468–488 ADD at cold and 512–564 ADD at warm)*; (ii)* T2 (end of the experiment): close to yolk sac resorption (740–752 ADD at cold and 862–866 ADD at warm). At both timepoints, all individuals were pooled by family and photographed using a digital camera (Nikon D5300, Nikon DX 18–105 mm lens). Measurements were taken using ImageJ software (v. 1.51, NIH, USA; Schneider *et al*., 2012). Length was measured as the total length (snout-tail length) in mm, and yolk sac volume (YSV) was calculated after considering yolk sac shape as an ellipsoid (Kamler, 2008), using the formula:

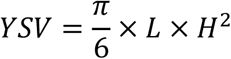

where L and H represent yolk sac length (longest axis) and yolk sac height/width (shorter axis, perpendicular to L axis), respectively (Figure S1).

Yolk sac conversion efficiency (YCE) was calculated at the family level as the ratio between somatic growth and yolk reserve depletion between T1 (hatching) and T2 (near yolk sac resorption), providing an estimate of the conversion of endogenous yolk reserves into somatic growth during early development. This metric follows the general concept of yolk conversion efficiency used in fish early-life studies (Kinnison *et al*., 1998; Fraser *et al*., 2010) and is expressed as the change in total length (mm) per change in yolk sac volume (mm^3^). Specifically, YCE (mm mm^-3^) was estimated as:

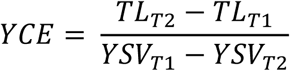

where TL represents total length (mm) and YSV represents yolk sac volume (mm^3^). Because individuals were pooled by family and individual estimates were not available, YCE was computed using mean values per family.

### Molecular analyses

Fin clips were collected from all adults and preserved in 95° ethanol. DNA was extracted using a Qiagen BioSprint 96 extraction robot. The extraction, PCR cycles, and multiplex genotyping were performed as indicated in Savary *et al*. (2017). We used seven neutral microsatellite markers known to be polymorphic in Arctic charr (Brunner *et al*., 1998; Savary *et al*., 2017).

To characterize population genetic structure and investigate potential admixture among populations given their respective stocking history, we used a Bayesian clustering approach implemented in STRUCTURE 2.3.4 (Pritchard *et al*., 2000). Analyses were conducted under the admixture model with a burn-in period of 100,000 iterations and 100,000 MCMC iterations. We performed runs for 1 to 5 clusters (K=1 to K=5) with 10 independent runs for each K to quantify the variation in likelihood, and calculated the logarithm of the data mean posterior probability LnP(K). We used STRUCTURE HARVESTER (Earl & vonHoldt, 2012) to summarize log-likelihood values across replicate runs and compute the Δ*K* statistic using Evanno’s method (Evanno *et al*., 2005) for identifying hierarchical levels of genetic structure. To account for label switching among replicate STRUCTURE runs, individual membership coefficients (Q-matrices) were aligned using CLUMPP (Jakobsson & Rosenberg, 2007) prior to visualization.

Neutral genetic differentiation was estimated using pairwise *F_st_* based on the method of Weir and Cockerham (1984) implemented in the R package *hierfstat* (Goudet, 2005). A 95% confidence interval (CI) for the observed *Fst* value was estimated using a standard bootstrap procedure with 10,000 iterations. To further visualize genetic relationships among populations, we performed a discriminant analysis of principal components (DAPC) using the R package *adegenet* (Jombart, 2008). Three principal components and three discriminant functions were retained, following the *k* − 1 criterion recommended for DAPC parameterization (Thia, 2023).

To assess within-population genetic diversity, we estimated the observed heterozygosity (*Ho*), expected heterozygosity (*He*), inbreeding coefficient (*Fis*) and allelic richness (*Ar*) using the functions *basic.stats* and *allelic.richness* implemented in the R package *hierfstat* (Goudet, 2005), with allele counts rarefied to the minimum sample size across populations (N = 20). The robustness of Ar estimates to sample size imbalance was further evaluated using 10,000 balanced random subsamplings (Leberg, 2002).

### Statistical analyses

To test the difference in thermal reaction norms among populations, we fitted a model per trait using linear mixed-effects models (LMMs) and generalized linear mixed models (GLMMs) using the functions *lmer* and *glmer* from the lme4 package (Bates *et al*., 2015). First, individual size at hatching (T1) and near yolk resorption (T2), yolk sac volume at T1 and T2, and incubation duration (in ADD) were set as response variable, and temperature, population and their interaction were included as fixed factors. Sire, dam and sire × dam interaction nested within population and block were fitted as random effects, together with plate for incubation duration and T1 traits, and incubator for T2 traits. Incubation duration (ADD) was analysed at the family x temperature level to avoid pseudo-replication, as ADD values were identical for all individuals within a given incubation group. Then, differences in thermal reaction norms on survival from eyed stage to T1 were analysed using GLMMs using a binomial error distribution and logit link function, with the survival status (0 = dead, 1 = alive) used as the response variable. Only plate was retained as a random effect because more complex random-effect structures did not converge reliably. Because hatching was asynchronous, individuals measured at T1 on the same calendar differed in thermal age. To account for this variation, we included ADD at hatching as a covariate in models of length and yolk sac volume at T1. Egg size is an important proxy for maternal effects, therefore, average egg size per female was added as a covariate in size, yolk sac, survival and incubation models (Kamler, 2005). Fixed effects were specified a priori and evaluated using likelihood ratio tests between hierarchically nested mixed-effects models.

Thermal reaction norms were visualized using estimated marginal means extracted from the fitted models with the *emmeans* package (Lenth *et al*., 2025). Traits were analysed at T1 for all populations in both treatments, and at T2 for all populations at warm and for three populations only at cold, due to an unexpected mortality event affecting Allos individuals during the cold treatment. Because of differences in spawning dates among populations, Allos was the only population being reared at that time.

### Genetic, phenotypic and quantitative differentiation analyses

To investigate whether phenotypic differentiation among populations are due to adaptive or neutral evolutionary processes, we compared neutral genetic differentiation (*F_st_*) with quantitative trait divergence (*Q_st_*), which assesses differentiation in quantitative traits (Spitze, 1993; Leinonen *et al*., 2013). For each trait, *Q_st_* was estimated both overall (among all populations) and pairwise (between population pairs) to assess differentiation among populations differing in origin (native or introduced) or habitat (low or high elevation). Within each temperature treatment, *Q_st_* was estimated for all traits using the following formula:

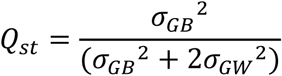

Where *σ_GB_*^2^ and *σ_GW_*^2^ represent the additive genetic variance components among and within populations, respectively (Merilä & Crnokrak, 2001; Leinonen *et al*., 2013). *σ_GB_*^2^ was estimated as the variance component among populations, and *σ_GW_*^2^ was estimated from sire variance components following our factorial half-sib design (*σ_GW_*^2^ = 4(*σ_sire_*^2^). To estimate variance components, we fitted LMMs for continuous traits and GLMMs for survival, separately for each temperature treatment (cold and warm). Random intercepts included experimental block nested within population, as well as sire, dam and their interaction (family), all nested within population and block. For incubation duration (in ADD), the dam and the family interaction were not included because ADD showed no within-family variation and was analysed at the family level to avoid pseudo-replication. To account for shared rearing conditions, we included incubator (for continuous traits) or plate (for survival) as additional random intercepts. Egg dry weight was included as a fixed covariate to control for maternal influences. Finally, we estimated narrow-sense heritability (ℎ^2^) as 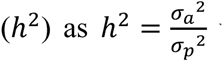 where 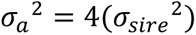 is the additive genetic variance and *σ_p_*^2^ = *σ_a_*^2^ + *σ_R_*^2^ is the total phenotypic variance within populations, with *σ_R_*^2^ denoting the residual variance (Lynch & Walsh, 1998). For early survival, fitted using binomial probit mixed models, heritability was estimated on the observed scale (i.e. the binary survival phenotype) using the *QGglmm* package (de Villemereuil, 2025). Confidence intervals around ℎ^2^ estimates were obtained using a parametric bootstrap procedure with 1,000 simulations implemented in the *bootMer* function of the *lme4* package (Bates *et al*., 2015). We assessed whether the observed *Q_st_* − *F_st_* significantly deviated from neutral expectations using the parametric approach described by Gilbert and Whitlock (2015). This approach compares the observed *Q_st_* − *F_st_* to a neutral distribution simulated from 10,000 bootstrap replicates based on neutral *F_st_* estimates and resampled genetic variance components.

All statistical analyses were performed using R (v. 4.4.2; R Core Team, 2024).

## Results

### Population genetic structure

Bayesian clustering analyses identified K = 2 as the most likely number of genetic clusters (Figure S1). Specifically, the introduced populations (Allos and Pavin) clustered primarily with their respective native source populations (Geneva and Constance, respectively), forming two genetic groups (Figure 2). Accordingly, pairwise *F_st_* estimates were lowest between Geneva and Allos (*F_st_* = 0.01; 95% CI: 0.00–0.02) and between Constance and Pavin (*F_st_* = 0.03; 95% CI: 0.01–0.05), whereas all other pairwise *F_st_* estimates ranged from 0.11 to 0.16 (Figure 2B). The overall *F_st_* among the four populations was 0.107 (95% CI: 0.074–0.138).

**Figure 2.**
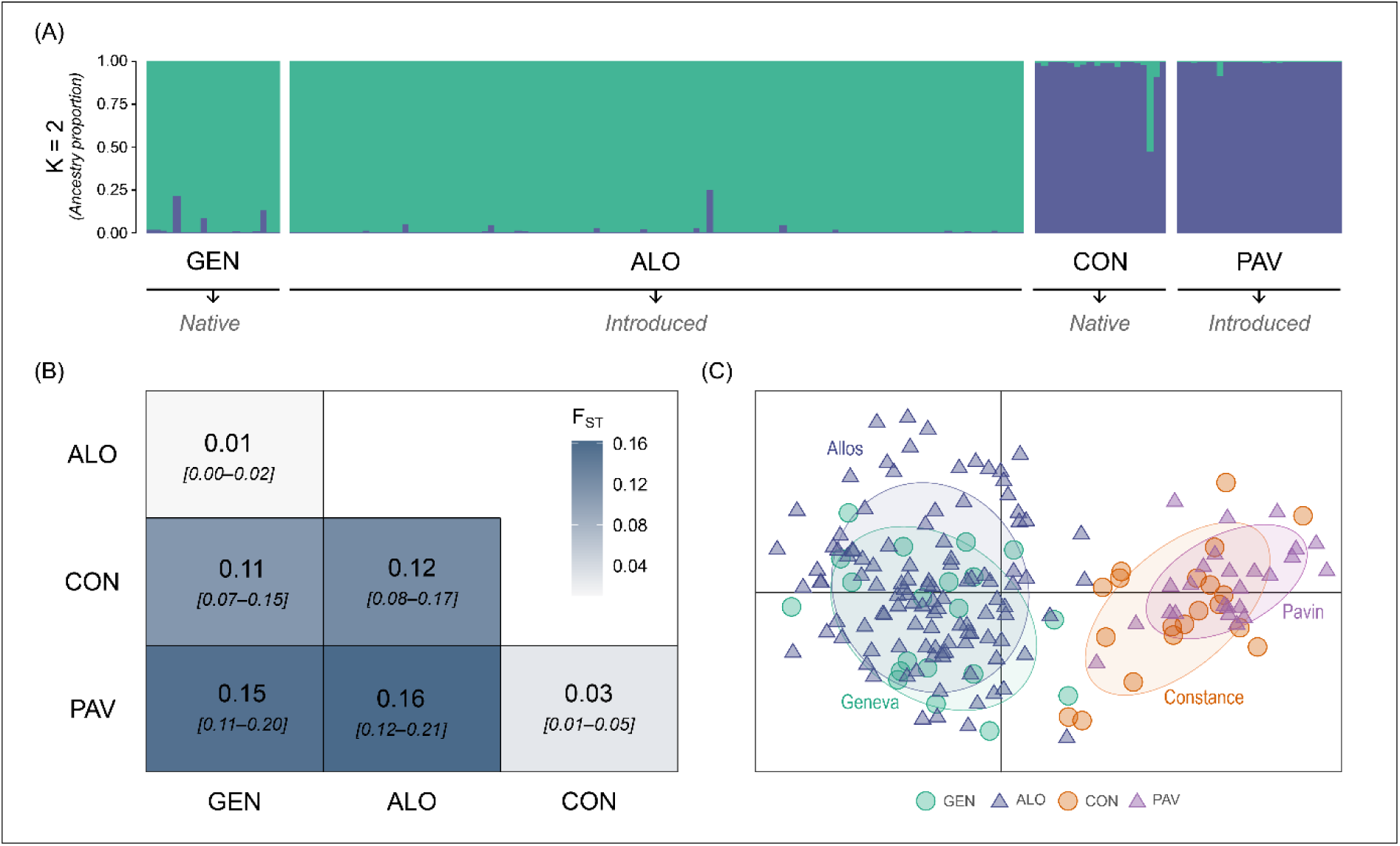
Genetic structure and differentiation among Arctic charr (*Salvelinus alpinus*) populations. A, Bayesian clustering analysis performed with STRUCTURE showing the ancestry proportions of individual genotypes for K = 2 clusters, grouped by population (Geneva, GEN, Allos, ALO, Constance, CON, Pavin, PAV). Each color corresponds to a distinct genetic cluster. B, pairwise *F_st_* estimates among four populations, calculated following Weir and Cockerham (1984). Values in brackets indicate 95% confidence intervals. All pairwise *F_st_* estimates were significant based on permutation tests, p < 0.05. C, Discriminant analysis of principal components (DAPC) showing genetic relationships among individuals. Ellipses represent 95% confidence intervals around population centroids. Circles and triangles indicate native and introduced populations, respectively.

Heterozygosity and allelic richness were higher in the Constance (*He* = 0.87*, Ar* = 11.86) and Pavin (*He* = 0.84*, Ar* = 9.74) populations than in the Geneva (*He* = 0.73*, Ar* = 7.43) and Allos (*He* = 0.74*, Ar* = 7.75) populations (Table S3). Population-level inbreeding coefficients (FIS) were positive across all populations and ranged from 0.13 to 0.20, with 95% confidence intervals overlapping zero only in the Pavin population (Table S3).

### Thermal reaction norms

Overall, all traits displayed significant thermal reaction norms, whose strength was population-specific (significant interaction between temperature and populations, Table 1, Figure 3). Warmer conditions significantly increased incubation duration in accumulated degree days (Figure 3A) and reduced survival at hatching across all populations (χ^2^ = 132, p < 0.001; Figure 3B). Incubation duration also differed among populations and exhibited a significant temperature x population interaction (χ^2^ = 212, p < 0.001). Across both temperature treatments, embryos from the Allos population hatched the earliest, whereas embryos from the Pavin population hatched the latest (Figure 3A).

**Figure 3.**
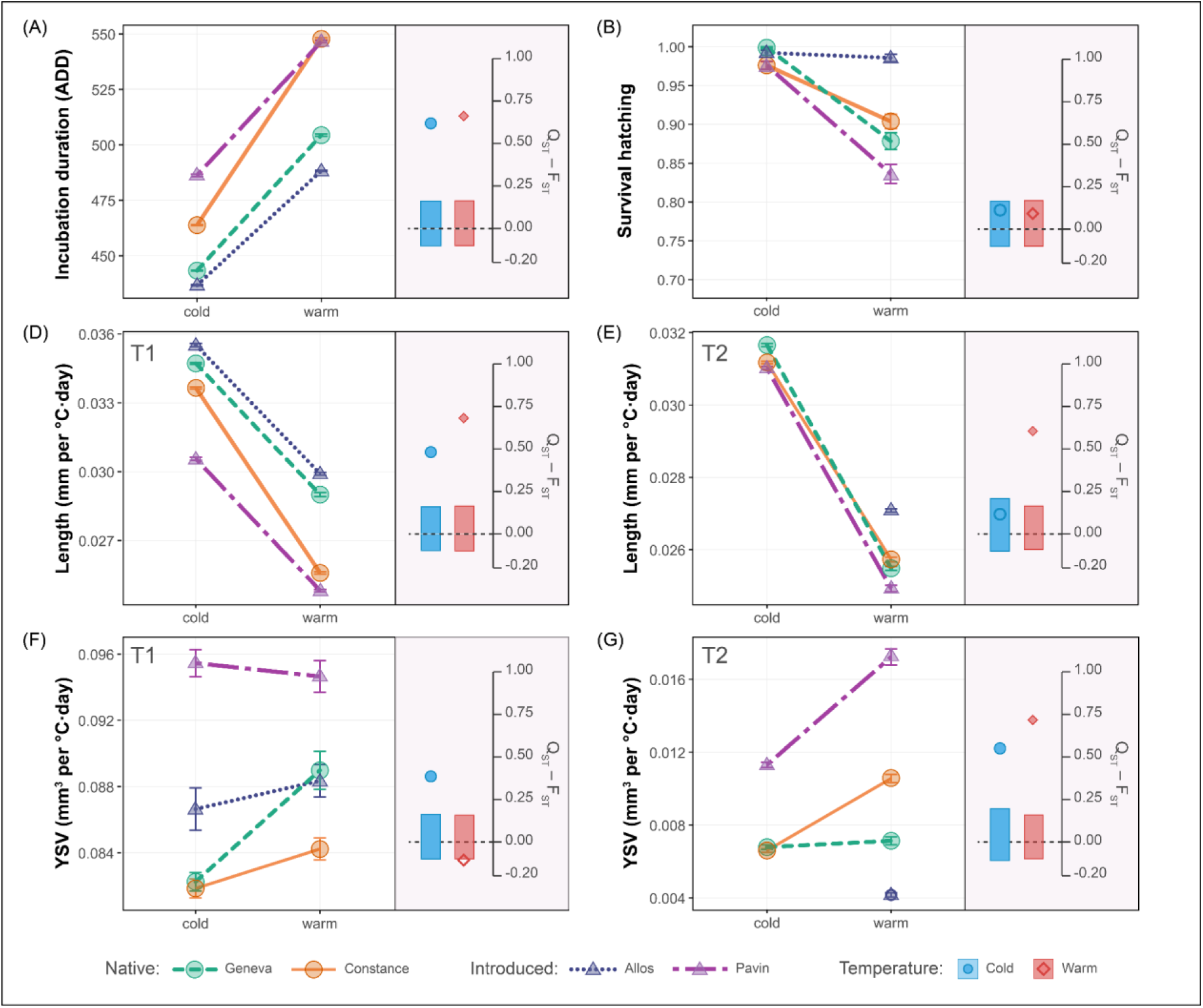
Thermal reaction norms and corresponding *Q_st_* − *F_st_* estimates for early life-history traits in four Arctic charr (*Salvelinus alpinus*) populations (Geneva, Allos, Constance, Pavin) reared under two temperature treatments (cold: 5 °C, warm: 8.5 °C). Left panels show thermal reaction norms for A, incubation duration in accumulated degree days (ADD), B, survival at hatching (proportion of successfully hatched individuals among those that reached the eyed stage), C, D, body length at hatching (T1, mm) and at the end of the experiment (T2, mm) and E, F, yolk sac volume (YSV, mm^3^) at T1 and T2. Circles indicate native populations from warm lakes (Geneva: green, Constance: orange), whereas triangles indicate populations introduced in cold lakes (Allos: blue, Pavin: purple). Values represent estimated marginal means adjusted for egg dry weight and ADD (where applicable) and error bars represent standard errors. Right panels display the corresponding *Q_st_* − *F_st_* values under cold (blue circles) and warm (orange diamonds) rearing conditions. Rectangles represent the simulated neutral expectation distribution of *Q_st_* − *F_st_*. Filled symbols indicate significant deviations from neutrality (p < 0.05), whereas open symbols indicate non-significant deviations.

**Table 1.** Effects of incubation temperature and population on early life-history traits.

| Fixed effects | *df* | $\chi^2$ | *p-value* | $\chi^2$ | *p-value* |
| --- | --- | --- | --- | --- | --- |
|  |  | Incubation duration (n = 443) |  | Survival at hatching (n = 6955) |  |
| Temperature | 1 | 944.88 | < 0.001 | 132.48 | < 0.001 |
| Population | 3 | 62.52 | < 0.001 | 57.49 | < 0.001 |
| T°C × population | 3 | 212.03 | < 0.001 | 26.10 | < 0.001 |
| egg dry weight | 1 | 0.35 | 0.55 | 4.87 | 0.03 |
|  |  | Length T1 (n = 4946) |  | Yolk sac volume T1 (n = 4945) |  |
| Temperature | 1 | 14.03 | < 0.001 | 0.42 | 0.52 |
| Population | 3 | 26.89 | < 0.001 | 7.01 | 0.07 |
| T°C × population | 3 | 103.81 | < 0.001 | 28.46 | < 0.001 |
| Egg dry weight | 1 | 32.67 | < 0.001 | 43.52 | < 0.001 |
| ADD | 1 | 27.16 | < 0.001 | 22.45 | < 0.001 |
|  |  | Length T2 (n = 5010) |  | Yolk sac volume T2 (n = 5010) |  |
| Temperature | 1 | 416.05 | < 0.001 | 202.73 | < 0.001 |
| Population | 3 | 62.10 | < 0.001 | 70.23 | < 0.001 |
| T°C × population | 2 | 12.88 | < 0.01 | 71.88 | < 0.001 |
| Egg dry weight | 1 | 38.10 | < 0.001 | 16.85 | < 0.001 |
|  |  | Yolk sac conversion efficiency (n = 224) |  |  |  |
| Temperature | 1 | 47.59 | < 0.001 |  |  |
| Population | 3 | 9.14 | 0.03 |  |  |
| T°C × population | 2 | 47.36 | < 0.001 |  |  |
| Egg dry weight | 1 | 37.87 | < 0.001 |  |  |

Likelihood ratio tests (LRT) from mixed-effects models testing the effects of incubation temperature (T°C), population, and their interaction (T°C × population) on incubation duration, survival at hatching, body length and yolk sac volume at hatching (T1) and at the end of the yolk sac stage (T2), and yolk sac conversion efficiency. Survival was analysed using binomial generalized linear mixed-effects models, whereas all other traits were analysed using linear mixed-effects models.

Survival at hatching was high under cold conditions in all populations but declined under warm conditions, with the decrease magnitude varying among populations, as indicated by the significant interaction between temperature and population (χ^2^ = 26.10, p < 0.001; Figure 3B). Overall, thermal reaction norms for survival differed among populations, with the lowest thermal sensitivity observed in Allos and the strongest decline in Pavin, while Geneva and Constance exhibited intermediate responses (Figure 3B).

Body length at hatching (T1) was significantly reduced at warm temperature (χ^2^ = 14.03, p < 0.001) and the intensity of this decrease differed among populations, with a significant temperature x population interaction (χ^2^ = 103.81, p < 0.001; Figure 3C). In contrast, yolk sac volume at hatching (T1) did not show an overall temperature effect but varied among populations through a significant interaction with temperature (χ^2^ = 28.46, p < 0.001; Figure 3E).

At the end of the yolk sac stage (T2), temperature had a strong negative effect on body length (χ^2^ = 416.05, p < 0.001), and population differences remained significant (χ^2^ = 62.10, p < 0.001), with significant temperature × population interaction (χ^2^ = 12.88, p < 0.01; Figure 3D). Yolk sac volume at T2 was strongly affected by temperature, population, and their interaction (all p < 0.001; Figure 3F). Offspring originating from Pavin displayed a flatter thermal reaction norm for yolk sac volume, whereas Geneva and Constance exhibited intermediate responses. Under warm conditions, individuals from Allos exhibited the lowest yolk sac volumes (Figure 3F).

Yolk sac conversion efficiency was affected by temperature (χ^2^ = 92.04, p < 0.001), and population differences remained significant (χ^2^ = 9.14, p = 0.03), with a significant temperature × population interaction (χ^2^ = 47.36, p < 0.001; Table 1). Egg dry weight had a significant negative effect on YCE (χ^2^ = 37.87, p < 0.001; Table 1). Under cold conditions, YCE did not differ among Constance, Geneva and Pavin (p ≥ 0.15). Under warm conditions, Geneva exhibited lower YCE than Constance (p = 0.01), Allos (p = 0.02) and Pavin (p < 0.001). Pavin showed higher YCE than Constance (p = 0.031). Other pairwise contrasts were not significant (Figure S3).

### Quantitative trait differentiation

Overall *Q_st_* ranged from 0 to 0.82 across traits and temperature treatments (Table S4; Figure 3). Comparisons between observed *Q_st_* − *F_st_* values and simulated neutral differentiation revealed trait and temperature specific differences from neutrality. At the overall population level, incubation duration showed significant quantitative differentiation exceeding neutral expectations under both thermal conditions (cold: *Q_st_*= 0.73, p < 0.001; warm: *Q_st_*= 0.77, p < 0.001). Body length at hatching (T1) exhibited significant levels of differentiation under both temperature treatments (cold: *Q_st_* = 0.59, p < 0.001; warm: *Q_st_* = 0.79, p < 0.001), indicating consistent divergence among populations. In contrast, survival at hatching showed no significant deviation from neutrality under either temperature (cold: *Q_st_* = 0.21, p = 0.07; warm: *Q_st_*= 0.20, p = 0.09). Patterns of differentiation differed at the end of the experiment (T2). Body length showed no significant differentiation under cold conditions (*Q_st_*= 0.21, p = 0.10) but exhibited significant divergence under warm conditions (*Q_st_*= 0.70, p < 0.001). Similarly, yolk sac volume at T2 displayed significant quantitative differentiation under warm (*Q_st_*= 0.82, p < 0.001), and cold conditions (*Q_st_*= 0.66, p < 0.001). Yolk sac conversion efficiency showed temperature-specific quantitative differentiation. Under cold conditions, the sire variance component was estimated at zero, which did not allow estimation of *Q_st_* (Table S4). Under warm conditions, YCE exhibited significant quantitative differentiation, exceeding neutral genetic expectations (*Q_st_* − *F_st_* = 0.59, p < 0.001; Table S4).

Pairwise *Q_st_* − *F_st_* comparisons between native populations and their introduced counterparts showed substantial heterogeneity among traits (Figure S4; Table S4). Several traits exhibited no detectable quantitative divergence, resulting in non-estimable or non-significant *Q_st_* values in multiple comparisons (survival at hatching and yolk sac volume at T1; Figure S3).

Narrow-sense heritability (*h^2^*) varied among traits, populations and temperature treatments (Table S4). Survival at hatching showed very low to near-zero *h^2^* overall, with near-zero estimates under cold conditions in most comparisons, except for Geneva–Allos (*h^2^* = 0.44), and low to near-zero values under warm conditions (*h^2^* = 0.04–0.12). Morphological traits showed low to moderate *h^2^* values (0.05–0.33), with generally lower estimates under warm conditions at hatching (T1), whereas estimates at T2 were more variable among population comparisons. In contrast, incubation duration displayed high *h^2^* under both cold (*h^2^* = 0.73-0.89) and warm conditions (*h^2^* = 0.75–0.89), with only minor reductions in some population combinations. For several traits by population combinations, additive genetic variance was close to zero, resulting in low or non-estimable *h^2^*values. For incubation duration and body length at hatching, additive genetic variance was generally lower in the Allos and Geneva populations compared to Pavin and Constance. Yolk sac conversion efficiency displayed narrow-sense heritability under warm conditions (*h^2^* = 0.12).

## Discussion

Phenotypic divergence can emerge early in development when populations experience contrasting thermal environments (Buckley & Kingsolver, 2021). By combining a common garden experimental design with quantitative genetic analyses, we quantified thermal reaction norms and divergence in early life-history traits among native and introduced Arctic charr populations originating from thermally contrasted habitats. Overall, temperature strongly affected early life-history traits, and reaction norms differed among populations in both magnitude and slope. We initially hypothesized that populations inhabiting warmer environments (e.g. Geneva and Constance) would exhibit higher performances (e.g., higher survival) under warm conditions than populations from colder habitats (e.g. Allos and Pavin), as predicted under local adaptation. However, this hypothesis was not supported by our results. In particular, the relatively high performance of the Allos population under elevated temperature, despite originating from the coldest environment included in this study, contradicts the expectation that contemporary habitat temperature directly predicts thermal performance. This result suggests that factors other than thermal conditions contribute to shape differences in thermal responses among populations. Together with the detection of additive genetic variance, these results also indicate that early developmental traits have the potential to respond to selection and may influence fitness through their effects on survival, growth, and recruitment (Elliott & Elliott, 2010; Massey & Hutchings, 2021; Jonsson *et al*., 2022).

### Population-specific thermal reaction norms

Contrary to our initial expectations, populations originating from warmer environments did not consistently exhibit higher performance under elevated temperature. In particular, the introduced Allos population exhibited higher survival and larger body size at T2 under warm conditions despite originating from the coldest environment. Conversely, the introduced Pavin population exhibited lower survival and body size than its native source population Constance under the same conditions (i.e. both warm and cold).

Accordingly, the thermal reaction norms differed in all populations, revealing *G* × *E* interactions for early life-history traits. Significant population × temperature interactions indicated that populations contrasted in their sensitivity to temperature, reflecting differences in both the magnitude and shape of plasticity responses across developmental stages. Such patterns are the phenotypic expression of *G* × *E* effects, i.e., genetic differences among populations in both baseline trait expression (intercepts) and plastic responses to warming (slopes), and are central to predicting evolutionary responses to climate change because selection can act on trait means, on plasticity itself, or on their covariance (Falconer, 1952; Via & Lande, 1985; Ghalambor *et al*., 2007; Saltz *et al*., 2018).

Higher temperature consistently resulted in shorter incubation time in calendar days but longer incubation duration in ADD, reduced body size, and bigger yolk sac volumes, indicating that individuals hatched at an earlier developmental stage compared to conspecifics reared under colder conditions (Kamler, 2008). Importantly, the magnitude of these effects differed strongly among populations. For instance, the survival of individuals from Lake Allos was less affected by higher temperature in contrast to other populations. This pattern suggests a reduced sensitivity of early survival to high temperatures. Moreover, population-specific plasticity in yolk sac volume suggested that some populations may partially buffer the negative effect of temperature (assuming that 5°C is a near-ideal temperature for the development of our Arctic charr populations).

Similarly, body length reaction norms diverged across populations at both hatching and near yolk sac resorption, while yolk sac volume responses differed markedly among populations. Together, these patterns indicate divergence in thermal sensitivity of energy allocation and growth efficiency during early development. Because body length and yolk utilization are tightly linked to fitness in salmonids, through recruitment success, size-mediated performance, and carry-over effects, divergence in reaction norms may have important fitness consequences (Pankhurst & Munday, 2011; Debes *et al*., 2021).

Rather than reflecting adaptation to present-day thermal regimes alone, the observed reaction norms may instead reflect differences in population history. Since populations living in warmer habitats did not display higher performance at warmer temperature, demographic fluctuations, introduction events, and supportive breeding practices may also contribute to shape the thermal responses observed among populations (Hutchings, 2011; Oomen & Hutchings, 2015; Solberg *et al*., 2016). Together, these results highlight differences among populations in trait expression under thermal stress, suggesting variation in early developmental trajectories and resource allocation strategies, as well as the role of reaction norms in shaping responses to environmental change (Pankhurst & Munday, 2011; Debes *et al*., 2021; Massey & Hutchings, 2021; Urban *et al*., 2024). Further study may aim at teasing apart the respective contributions of these processes to better understand responses to thermal stress.

### Neutral and quantitative population divergence

Pairwise *Q_st_* − *F_st_* comparisons revealed pronounced heterogeneity among population pairs. Strong positive deviations from neutrality were most frequently observed for body size traits, particularly in comparisons involving the Pavin population. In contrast, several traits, especially survival at hatching and yolk sac volume at T1, showed no detectable between-population genetic variance in multiple pairwise comparisons. Comparisons between the two native populations, Geneva and Constance showed quantitative divergence for several traits and temperatures. This is in accordance with long term evolutionary divergence, but more contemporary demographic events and practices (stocking including with non native populatoins or fishing) could also lead to such pattern and their effect can not be ruled out. However, for some traits, such as yolk sac volume, observed *Q_st_* − *F_st_* deviations may partly reflect maternal contributions in addition to additive genetic differentiation, given the maternal effects on this trait in salmonids (Janhunen *et al*., 2010; Leblanc *et al*., 2016). Comparisons between source lakes and their introduced populations (Geneva–Allos and Constance–Pavin) showed positive *Q_st_* − *F_st_* deviations for incubation duration and body length in both population pairs, as well as yolk sac volume at T1 in the Geneva–Allos comparison and at T2 in the Constance–Pavin comparison. Quantitative divergence therefore did not scale proportionally with neutral differentiation and may reflect rapid evolutionary processes following population establishment (Rogissart *et al*., 2026). Because these introduced populations were established less than 150 years ago, the repeated *Q_st_* − *F_st_* deviations observed for several traits suggest that adaptive divergence may occur over relatively short timescales (Koskinen *et al*., 2002; Rogissart *et al*., 2026).

### Within-population genetic variation

Additive genetic variance showed substantial variation among traits, populations and temperature treatments. Heritability estimates were particularly low for survival at hatching. This is consistent with expectations that traits under strong directional selection tend to exhibit reduced additive genetic variance (Merilä & Sheldon, 1999). More generally, heritability estimates tended to be lower under warm conditions for most traits, consistent with previous findings that environmental stress can modify the expression of quantitative genetic variation and influence heritability estimates (Charmantier & Garant, 2005). However, the magnitude of this reduction differed among traits and populations, suggesting that genetic variance does not respond uniformly to thermal stress (Urban *et al*., 2024). For morphological traits at hatching (T1), *h^2^* was generally lower under warm conditions, whereas estimates at later stages (T2) were more variable across population comparisons. In contrast, incubation duration maintained consistently high *h^2^* across temperatures, with only moderate reductions in some cases. Together, these results indicate that temperature did not cause a uniform reduction in genetic variation. Differences among populations may reflect the combined effects of contrasting environments, demographic history, introduction events, and management practices. However, given the limited number of populations examined and the absence of replicated management histories, our data do not allow us to disentangle the specific effects of ecological and evolutionary processes from stocking. At the very least, it seems clear that management history complicates both the scientific analysis of evolutionary processes and the potential of populations to respond to environmental change (Ford, 2002; Christie *et al*., 2012).

## Conclusion

We expected higher temperatures to have drastic effects on early life-history traits and reveal adaptive differences in favor of populations from warmer environments. Our results do not support this prediction and instead show that responses to temperature are population- and trait-specific. We detected genetic population divergence in early life-history traits, consistent with selection driven differentiation among contrasting environments, although factors other than temperature may also contribute to this divergence. These results further suggest that adaptive divergence can occur over relatively short timescales, as illustrated by the differentiation detected between native and recently introduced populations established less than approximately 150 years ago. Future research integrating thermal reaction norms across early life stages and broad temperature ranges, together with transcriptomic approaches investigating the physiological mechanisms underlying thermal responses, will improve our understanding of population resilience to climate warming.

## Acknowledgements

We acknowledge the APERA (Association pour la mise en valeur piscicole des Plans d’Eaux en Rhône-Alpes), the Mercantour National Park, LAZBW (Landwirtschaftliches Zentrum für Rinderhaltung, Grünlandwirtschaft, Milchwirtschaft, Wild und Fischerei Baden-Württemberg) and the Fédération de Pêche du Puy de Dôme staff for their contribution of Arctic charr spawners, granting fishing permissions and logistical assistance. We thank Laurent Espinat and Ayala Loisel for their assistance during field sampling and Céline Stoffel for technical help for genotyping analyses; and the Environmental Physics group at the Limnological Institute of the University of Konstanz for providing temperature data for Lake Constance.

## CRediT Statement

Hervé Rogissart (Methodology, Data curation, Formal analysis, Validation, Visualization, Writing – original draft, Writing – review & editing), Lisandrina Mari (Conceptualization, Methodology, Data curation, Formal analysis, Validation, Writing – original draft, Writing – review & editing), Guillaume Evanno (Conceptualization, Methodology, Writing – review & editing), Martin Daufresne (Conceptualization, Methodology, Writing – review & editing), Luca Fumagalli (Supervision, Writing – review & editing), Jean Guillard (Writing – review & editing), Allan Raffard (Writing – review & editing), and Emilien Lasne (Funding acquisition, Project administration, Supervision, Conceptualization, Methodology, Writing – review & editing).

## Declaration of Competing Interest

The authors declare that they have no known competing financial interests or personal relationships that could have appeared to influence the work reported in this paper.

## Funding

Funding for field and experimental work was provided by Office Français de la Biodiversité (OFB). HR was funded by the CSMB (Conseil Savoie Mont Blanc) and AQUA department (INRAE). LM was funded by the French Ministry for Research.

## Data Availability Statement

The data supporting the findings of this study are available on Figshare.

## Ethics

This research was designed in accordance with European Directive 2010/63/EU. As this study focused on fish embryos and larvae that rely on endogenous feeding only, no special ethics permission was required. Permission for fish captures on the field was granted by local authorities.

## SUPPORTING INFORMATION

**Table S1.** Main geographical and historical characteristics of the four study lakes and their Arctic charr (*Salvelinus alpinus*) populations.

**Table S2.** Crossing design used for the experimental breeding of four Arctic charr (*Salvelinus alpinus*) populations, based on wild-caught spawners.

**Figure S1.** Morphometric measurements of Arctic charr (*Salvelinus alpinus*) larvae at hatching used to estimate yolk sac volume (YSV).

**Figure S2.** Determination of the number of genetic clusters and population genetic structure in Arctic charr (*Salvelinus alpinus*).

**Table S3.** Neutral genetic diversity at seven microsatellite loci across four Arctic charr (*Salvelinus alpinus*) populations.

**Figure S3.** Thermal reaction norms and *Q_st_* − *F_st_* estimates for yolk sac conversion efficiency (YCE) in four Arctic charr populations reared under cold (5 °C) and warm (8.5 °C) conditions.

**Figure S4.** Simulated neutral expectations and observed values of *Q_st_* − *F_st_* for population comparisons in Arctic charr (*Salvelinus alpinus*).

**Table S4.** Summary of the linear mixed models and generalized linear mixed-effects models assessing the variance components and *Q_st_* estimates for quantitative traits and early survival in *Salvelinus alpinus*.

## Notes

### Competing Interest Statement

The authors have declared no competing interest.

### Summary of Updates

You are viewing Version 2, the most recent version of this article.

## References

1. Alexander, T. & Seehausen, O. 2021. *Diversity, distribution and community composition of fish in perialpine lakes. “Projet Lac” synthesis report*. Swiss Federal Institute of Aquatic Science and Technology, Eawag.

2. Angilletta, M.J., Niewiarowski, P.H. & Navas, C.A. 2002. The evolution of thermal physiology in ectotherms. Journal of Thermal Biology 27: 249–268.

3. Barley, J.M., Cheng, B.S., Sasaki, M., Gignoux-Wolfsohn, S., Hays, C.G., Putnam, A.B., et al. 2021. Limited plasticity in thermally tolerant ectotherm populations: evidence for a trade-off. Proc. R. Soc. B. 288: 20210765.

4. Bates, D., Mächler, M., Bolker, B. & Walker, S. 2015. Fitting Linear Mixed-Effects Models Using lme 4. J. Stat. Soft. 67.

5. Bebak, J., Hankins, J.A. & Summerfelt, S.T. 2000. Effect of Water Temperature on Survival of Eyed Eggs and Alevins of Arctic Char. North American Journal of Aquaculture 62: 139–143.

6. Blaxter, J.H.S. 1991. The Effect of Temperature On Larval Fishes. Neth J Zool 42: 336–357.

7. Bogataj, L.K. 2007. How will the Alps respond to climate change? Scenarios for the future of alpine water. Alpine Space: Man and Environment 3: 43–51.

8. Brunner, P.C., Douglas, M.R. & Bernatchez, L. 1998. Microsatellite and mitochondrial DNA assessment of population structure and stocking effects in Arctic charr Salvelinus alpinus (Teleostei: Salmonidae) from central Alpine lakes. Molecular Ecology 7: 209–223.

9. Brunner, P.C., Douglas, M.R., Osinov, A., Wilson, C.C. & Bernatchez, L. 2001. Holarctic phylogeography of Arctic charr (Salvelinus alpinus L.) inferred from mitochondrial DNA sequences. Evolution 55: 573–586.

10. Buckley, L.B. & Kingsolver, J.G. 2021. Evolution of Thermal Sensitivity in Changing and Variable Climates. Annu. Rev. Ecol. Evol. Syst. 52: 563–586.

11. Caudron, A., Lasne, E., Gillet, C., Guillard, J. & Champigneulle, A. 2014. Thirty years of reoligotrophication do not contribute to restore self-sustaining fisheries of Arctic charr, *Salvelinus alpinus*, in Lake Geneva. Fisheries Research 154: 165–171.

12. Charmantier, A. & Garant, D. 2005. Environmental quality and evolutionary potential: lessons from wild populations. Proc. R. Soc. B. 272: 1415–1425.

13. Chevin, L.-M. & Bridle, J. 2025. Impacts of limits to adaptation on population and community persistence in a changing environment. Phil. Trans. R. Soc. B 380: 20230322.

14. Christie, M.R., Ford, M.J. & Blouin, M.S. 2014. On the reproductive success of early-generation hatchery fish in the wild. Evolutionary Applications 7: 883–896.

15. Christie, M.R., Marine, M.L., French, R.A. & Blouin, M.S. 2012. Genetic adaptation to captivity can occur in a single generation. Proc. Natl. Acad. Sci. U.S.A. 109: 238–242.

16. Coumes, J. 1862. *N*otice historique sur l’établissement de pisciculture de Huningue. Veuve Berger-Levrault, Strasbourg.

17. Coutant, C. 1976. Thermal effects on fish ecology. In: Encyclopedia of Environmental Science and Engineering, pp. 891–896. Gordon and Breach Publishers, New York, NY.

18. Crettiez, J.M. 1906. De la culture de l’omble-chevalier du lac Léman métis et hybrides de ce salmonide. In: Compte rendu de l’Association Française pour l’Avancement des Sciences, section Zoologie, pp. 498–506. Lyon.

19. Dahlke, F., Butzin, M., Wohlrab, S. & Pörtner, H.-O. 2022. Reply to: methodological inconsistencies define thermal bottlenecks in fish life cycle. Evol Ecol 36: 293–298.

20. Dahlke, F.T., Wohlrab, S., Butzin, M. & Pörtner, H.-O. 2020. Thermal bottlenecks in the life cycle define climate vulnerability of fish. Science 369: 65–70.

21. Dayan, D.I., Sard, N.M., Johnson, M.A., Fitzpatrick, C.K., Couture, R. & O’Malley, K.G. 2024. A single generation in the wild increases fitness for descendants of hatchery-origin Chinook salmon (*Oncorhynchus tshawytscha*). Evolutionary Applications 17: e13678.

22. De March, B.G.E. 1995. Effects of Incubation Temperature on the Hatching Success of Arctic Char Eggs. The Progressive Fish-Culturist 57: 132–136.

23. de Villemereuil, P. 2025. How to use the QGglmm package?

24. Debes, V., Paul, Solberg, M.F., Matre, I.H., Dyrhovden, L. & Glover, K.A. 2021. Genetic variation for upper thermal tolerance diminishes within and between populations with increasing acclimation temperature in Atlantic salmon. Heredity 127: 455–466.

25. Desmolles, F. 2016. History of the Fish Fauna of Lake Pavin: A Population Heavily Influenced by Man? In: Lake Pavin (T. Sime-Ngando, P. Boivin, E. Chapron, D. Jezequel, & M. Meybeck, eds), pp. 345–361. Springer International Publishing, Cham.

26. Dumont, M., Monteiro, D., Filhol, S., Gascoin, S., Marty, C., Hagenmuller, P., et al. 2025. The European Alps in a changing climate: physical trends and impacts. Comptes Rendus. Géoscience 357: 25–42.

27. Dussart, B. 1955. Un Poisson trop peu connu en France, l’omble chevalier. revec 9: 129–158.

28. Earl, D.A. & vonHoldt, B.M. 2012. STRUCTURE HARVESTER: a website and program for visualizing STRUCTURE output and implementing the Evanno method. Conservation Genet Resour 4: 359–361. Springer Science and Business Media LLC.

29. Elliott, J.M. & Elliott, J.A. 2010. Temperature requirements of Atlantic salmon *Salmo salar*, brown trout *Salmo trutta* and Arctic charr *Salvelinus alpinus* : predicting the effects of climate change. Journal of Fish Biology 77: 1793–1817.

30. Evanno, G., Regnaut, S. & Goudet, J. 2005. Detecting the number of clusters of individuals using the software STRUCTURE: a simulation study. Molecular Ecology 14: 2611–2620. Wiley.

31. Falconer, D.S. 1952. The Problem of Environment and Selection. The American Naturalist 86: 293– 298.

32. Ford, M.J. 2002. Selection in Captivity during Supportive Breeding May Reduce Fitness in the Wild. Conservation Biology 16: 815–825.

33. Fraser, D.J., Houde, A.L.S., Debes, P.V., O’Reilly, P., Eddington, J.D. & Hutchings, J.A. 2010. Consequences of farmed–wild hybridization across divergent wild populations and multiple traits in salmon. Ecological Applications 20: 935–953.

34. Gendron, H.K., Tallman, R.F. & Docker, M.F. 2025. The influence of climate change on growth of Arctic charr (*Salvelinus alpinus*). Environ Biol Fish 108: 2087–2104.

35. Ghalambor, C.K., McKAY, J.K., Carroll, S.P. & Reznick, D.N. 2007. Adaptive versus non-adaptive phenotypic plasticity and the potential for contemporary adaptation in new environments. Functional Ecology 21: 394–407.

36. Gienapp, P., Teplitsky, C., Alho, J.S., Mills, J.A. & Merilä, J. 2008. Climate change and evolution : disentangling environmental and genetic responses. Molecular Ecology 167–178.

37. Goudet, J. 2005. HIERFSTAT, a package for R to compute and test hierarchical *F* -statistics. Molecular Ecology Notes 5: 184–186.

38. Guillard, J., Gillet, C. & Champigneulle, A. 1992. Revue bibliographique - Principales caractéristiques de l’élevage de l’omble chevalier (*Salvelinus alpinus* L.) en eau douce. Bull. Fr. Pêche Piscic. 47–68.

39. Hemmer-Brepson, C., Replumaz, L., Romestaing, C., Voituron, Y. & Daufresne, M. 2013. Non-stressful temperature effect on oxidative balance and life history traits in adults fish (*Oryzias latipes*). Journal of Experimental Biology jeb.096172.

40. Hendry, A.P., Kinnison, M.T., Heino, M., Day, T., Smith, T.B., Fitt, G., et al. 2011. Evolutionary principles and their practical application. Evolutionary Applications 4: 159–183.

41. Hutchings, J.A. 2011. Old wine in new bottles: reaction norms in salmonid fishes. Heredity 106: 421– 437.

42. Jakobsson, M. & Rosenberg, N.A. 2007. CLUMPP: a cluster matching and permutation program for dealing with label switching and multimodality in analysis of population structure. Bioinformatics 23: 1801–1806. Oxford University Press.

43. Janhunen, M., Piironen, J. & Peuhkuri, N. 2010. Parental effects on embryonic viability and growth in Arctic charr *Salvelinus alpinus* at two incubation temperatures. Journal of Fish Biology 76: 2558–2570.

44. Jeuthe, H., Brännäs, E. & Nilsson, J. 2016. Effects of variable egg incubation temperatures on the embryonic development in Arctic charr *Salvelinus alpinus*. Aquac Res 47: 3753–3764.

45. Jombart, T. 2008. *adegenet* : a R package for the multivariate analysis of genetic markers. Bioinformatics 24: 1403–1405.

46. Jonsson, B. 2023. Thermal Effects on Ecological Traits of Salmonids. Fishes 8: 337.

47. Jonsson, B., Jonsson, N. & Hansen, M.M. 2022. Knock-On Effects of Environmental Influences during Embryonic Development of Ectothermic Vertebrates. The Quarterly Review of Biology 97: 95–139.

48. Jungwirth, M. & Winkler, H. 1984. The temperature dependence of embryonic development of grayling (*Thymallus thymallus*), Danube salmon (*Hucho hucho*), Arctic char (*Salvelinus alpinus*) and brown trout (*Salmo trutta fario*). Aquaculture 38: 315–327.

49. Kamler, E. 2005. Parent–egg–progeny Relationships in Teleost Fishes: An Energetics Perspective. Reviews in Fish Biology and Fisheries 15: 399–421.

50. Kamler, E. 2008. Resource allocation in yolk-feeding fish. Rev Fish Biol Fisheries 18: 143–200.

51. Kavanagh, K.D., Haugen, T.O., Gregersen, F., Jernvall, J. & Vøllestad, L.A. 2010. Contemporary temperature-driven divergence in a Nordic freshwater fish under conditions commonly thought to hinder adaptation. BMC Evol Biol 10: 350.

52. Kawecki, T.J. & Ebert, D. 2004. Conceptual issues in local adaptation. Ecology Letters 7: 1225–1241.

53. Kelly, S., Moore, T.N., De Eyto, E., Dillane, M., Goulon, C., Guillard, J., et al. 2020. Warming winters threaten peripheral Arctic charr populations of Europe. Climatic Change 163: 599–618.

54. King, H.R., Pankhurst, N.W. & Watts, M. 2007. Reproductive sensitivity to elevated water temperatures in female Atlantic salmon is heightened at certain stages of vitellogenesis. Journal of Fish Biology 70: 190–205.

55. Kinnison, M.T., Unwin, M.J., Hershberger, W.K. & Quinn, T.P. 1998. Egg size, fecundity, and development rate of two introduced New Zealand chinook salmon (*Oncorhynchus tshawytscha*) populations. Canadian Journal of Fisheries and Aquatic Sciences 55: 1946– 1953.

56. Klemetsen, A. 2013. The most variable vertebrate on Earth. J. Ichthyol. 53: 781–791.

57. Koskinen, M.T., Haugen, T.O. & Primmer, C.R. 2002. Contemporary fisherian life-history evolution in small salmonid populations. Nature 419: 826–830.

58. Larsson, S. & Berglund, I. 2005. The effect of temperature on the energetic growth efficiency of Arctic charr (*Salvelinus alpinus* L.) from four Swedish populations. Journal of Thermal Biology 30: 29–36.

59. Leberg, P.L. 2002. Estimating allelic richness: Effects of sample size and bottlenecks. Molecular Ecology 11: 2445–2449.

60. Leblanc, C.A.-L., Kristjánsson, B.K. & Skúlason, S. 2016. The importance of egg size and egg energy density for early size patterns and performance of Arctic charr *Salvelinus alpinus*. Aquac Res 47: 1100–1111.

61. Lehtonen, H. 1998. Does global warming threat the existence of Arctic charr, *Salvelinus alpinus* (Salmonidae), in northern Finland? Italian Journal of Zoology 65: 471–474.

62. Leinonen, T., McCairns, R.J.S., O’Hara, R.B. & Merilä, J. 2013. QST–FST comparisons: evolutionary and ecological insights from genomic heterogeneity. Nat Rev Genet 14: 179–190.

63. Lenth, R.V., Banfai, B., Bolker, B., Buerkner, P., Giné-Vázquez, I., Herve, M., et al. 2025. emmeans: Estimated marginal means, aka least-squares means.

64. Lynch, M. & Walsh, B. 1998. *Genetics and analysis of quantitative traits*. Sinauer Sunderland, MA.

65. Machino, Y. 1991. The Arctic charr (*Salvelinus alpinus*) in France: present geographical distribution of original and transplanted populations. Int. Soc. Arctic Charr Fanatics Inf. Ser. 5: 113–120.

66. Machino, Y. & Rivier, B. 2002. Le lac d’Allos (Alpes-de-Haute-Provence). Sanctuaire et laboratoire de l’évolution pour l’omble chevalier du Léman (*Salvelinus alpinus*, Salmonidae). Cybium 26: 173–177.

67. Mari, L., Daufresne, M., Guillard, J., Evanno, G. & Lasne, E. 2021. Elevated temperature and deposited sediment jointly affect early life history traits in southernmost Arctic char populations. Can. J. Fish. Aquat. Sci. 78: 744–751.

68. Mari, L., Garaud, L., Evanno, G. & Lasne, E. 2016. Higher temperature exacerbates the impact of sediments on embryo performances in a salmonid. Biol. Lett. 12: 20160745.

69. Massey, M.D. & Hutchings, J.A. 2021. Thermal variability during ectotherm egg incubation: A synthesis and framework. J Exp Zool Pt A 335: 59–71.

70. Merilä, J. & Crnokrak, P. 2001. Comparison of genetic differentiation at marker loci and quantitative traits. Journal of Evolutionary Biology 14: 892–903.

71. Merilä, J. & Sheldon, B.C. 1999. Genetic architecture of fitness and nonfitness traits: empirical patterns and development of ideas. Heredity 83: 103–109.

72. Muhlfeld, C.C., Cline, T.J., Finstad, A.G., Hessen, D.O., Perrin, S., Thaulow, J., et al. 2024. Climate change vulnerability of Arctic char across Scandinavia. Global Change Biology 30: e17387.

73. Muhlfeld, C.C., Dauwalter, D.C., D’Angelo, V.S., Ferguson, A., Giersch, J.J., Impson, D., et al. 2019. Chapter 21: Global status of trout and char: Conservation challenges in the twenty-first century. In: Trout and char of the world, pp. 717–760. American Fisheries Society, Bethesda, Maryland.

74. Narum, S.R., Campbell, N.R., Meyer, K.A., Miller, M.R. & Hardy, R.W. 2013. Thermal adaptation and acclimation of ectotherms from differing aquatic climates. Molecular Ecology 22: 3090– 3097.

75. O’Dea, R.E., Lagisz, M., Hendry, A.P. & Nakagawa, S. 2019. Developmental temperature affects phenotypic means and variability: A meta-analysis of fish data. Fish and Fisheries 20: 1005– 1022.

76. Olk, T.R., Jeuthe, H., Thorarensen, H., Wollebæk, J. & Lydersen, E. 2020. Brood-stock management and early hatchery rearing of Arctic charr (*Salvelinus alpinus* (Linnaeus)). Reviews in Aquaculture 12: 1595–1623.

77. Oomen, R.A. & Hutchings, J.A. 2015. Genetic variability in reaction norms in fishes. Environ. Rev. 23: 353–366.

78. Pankhurst, N.W. & King, H.R. 2010. Temperature and salmonid reproduction: implications for aquaculture. Journal of Fish Biology 76: 69–85.

79. Pankhurst, N.W. & Munday, P.L. 2011. Effects of climate change on fish reproduction and early life history stages. Marine and Freshwater Research 62: 1015–1026.

80. Pörtner, H.O. & Farrell, A.P. 2008. Physiology and Climate Change. Science 322: 690–692.

81. Pritchard, J.K., Stephens, M. & Donnelly, P. 2000. Inference of Population Structure Using Multilocus Genotype Data. Genetics 155: 945–959. Oxford University Press (OUP).

82. R Core Team. 2024. R: A language and environment for statistical computing. Vienna, Austria.

83. Realis-Doyelle, E., Gisbert, E., Alcaraz, C., Teletchea, F. & Pasquet, A. 2018. Temperature affects growth allometry and development patterns in brown trout (Salmo trutta) fry: a multitrait approach. Canadian Journal of Fisheries and Aquatic Sciences 75: 714–722.

84. Reynouard, J. 1909. Besse aujourd’hui autrefois. Annales de la station limnologique de Besse. In: Internal publication of Besse limnological and biological station, pp. 6–11. Clermont-Ferrand, France.

85. Rogissart, H., Daufresne, M., Evanno, G., Guillard, J., Lubin, F.-R., Chancerel, E., et al. 2026. A century of allopatry: plasticity and rapid selection shape phenotypic trait variability under contrasting environments. Proc. R. Soc. B 293: 20260692.

86. Sæther, B.-S., Siikavuopio, S.I. & Jobling, M. 2016. Environmental conditions required for intensive farming of Arctic charr (*Salvelinus alpinus* (L.)). Hydrobiologia 783: 347–359.

87. Saltz, J.B., Bell, A.M., Flint, J., Gomulkiewicz, R., Hughes, K.A. & Keagy, J. 2018. Why does the magnitude of genotype-by-environment interaction vary? Ecology and Evolution 8: 6342– 6353.

88. Savary, R., Dufresnes, C., Champigneulle, A., Caudron, A., Dubey, S., Perrin, N., et al. 2017. Stocking activities for the Arctic charr in Lake Geneva: Genetic effects in space and time. Ecology and Evolution 7: 5201–5211.

89. Schneider, C.A., Rasband, W.S. & Eliceiri, K.W. 2012. NIH Image to ImageJ: 25 years of image analysis. Nat Methods 9: 671–675.

90. Solberg, M.F., Dyrhovden, L., Matre, I.H. & Glover, K.A. 2016. Thermal plasticity in farmed, wild and hybrid Atlantic salmon during early development: has domestication caused divergence in low temperature tolerance? BMC Evol Biol 16: 38.

91. Spitze, K. 1993. Population structure in Daphnia obtusa: quantitative genetic and allozymic variation. Genetics 135: 367–374.

92. Stearns, S.C. 1992. Stearns, Stephen C., 1992. The Evolution of Life Histories. Oxford University Press, London xii + 249 pp. John Wiley & Sons, Ltd (10.1111).

93. Swift, D.R. 1965. Effect of temperature on mortality and rate of development of the eggs of the windermere char (*Salvelinus alpinus*). J. Fish. Res. Bd. Can. 22: 913–917.

94. Takatsu, K., Selz, O.M. & Brodersen, J. 2022. Temperature regime during embryogenesis alters subsequent behavioural phenotypes of juvenile brown trout. Biol. Lett. 18: 20220369.

95. Thia, J.A. 2023. Guidelines for standardizing the application of discriminant analysis of principal components to genotype data. Molecular Ecology Resources 23: 523–538. Wiley.

96. Urban, M.C., Swaegers, J., Stoks, R., Snook, R.R., Otto, S.P., Noble, D.W.A., et al. 2024. When and how can we predict adaptive responses to climate change? Evolution Letters 8: 172–187.

97. Via, S. & Lande, R. 1985. Genotype-environment interaction and the evolution of phenotypic plasticity. Evolution 39: 505–522.

98. Weinstein, S.Y., Gallagher, C.P., Hale, M.C., Loewen, T.N., Power, M., Reist, J.D., et al. 2024. An updated review of the post-glacial history, ecology, and diversity of Arctic char (*Salvelinus alpinus*) and Dolly Varden (*S. malma*). Environ Biol Fish 107: 121–154.

99. Weir, B.S. & Cockerham, C.C. 1984. Estimating *F*-Statistics for the Analysis of Population-Structure. Evolution 38: 1358–1370.

100. Williams, J.E., Kershner, J.L. & Zablocki, J.A. 2024. The Future of Salmonids in a Rapidly Changing World. In: Advances in the Ecology of Stream-Dwelling Salmonids (J. Lobon-Cervia, P. Budy, & R. Gresswell, eds), pp. 709–731. Springer International Publishing, Cham.

