## Supplementary materials for "Early life thermal plasticity and adaptive divergence among populations of Arctic charr (*Salvelinus alpinus*)"

**Table S1.** Main geographical and historical characteristics of the four study lakes and their Arctic charr (*Salvelinus alpinus*) populations.

| | *Native populations* | | *Introduced populations* | |
|---|---|---|---|---|
| **Parameter** | **Geneva** | **Constance** | **Allos** | **Pavin** |
| **Latitude (°N)** | 46°26'37" | 47°36'04" | 44°14'02" | 45°29'44" |
| **Longitude (°E)** | 6°31'15" | 9°24'51" | 6°42'29" | 2°53'17" |
| **Elevation (m a.s.l.)**[a] | 372 | 395 | 2232 | 1197 |
| **Surface area ($km^2$)** | 580 | 500 | 0.54 | 0.44 |
| **Max. depth (m)** | 309 | 252 | 51 | 92 |
| **Mean depth (m)** | 153 | 90 | 25 | 29.5 |
| **Founding date** | – | – | 1922, 1924 | 1859, 1900–1906 |
| **Founders (n)** | – | – | 2000 | 8000, 2500 |
| **Historic origin** | – | – | Geneva | Constance, Geneva |
| **Spawning grounds depth (m)** | 35–50 | 40–80 | 10–15 | 15 |

[a] Elevation is given in meters above sea level (m a.s.l.).

**Table S2.** Crossing design used for the experimental breeding of four Arctic charr (*Salvelinus alpinus*) populations, based on wild-caught spawners.

| Population | Dam (TL, mm) | Crossed sires (TL, mm) | Block |
|---|---|---|---|
| ***Native populations*** | | | |
| **Geneva** | F30 (437) | M40 (388), M41 (311), M42 (318), M43 (280) | G1 |
| | F31 (380) | M40 (388), M41 (311), M42 (318), M43 (280) | G1 |
| | F32 (430) | M40 (388), M41 (311), M42 (318), M43 (280) | G1 |
| | F33 (352) | M44 (373), M45 (353), M46 (347), M47 (332) | G2 |
| | F35 (443) | M44 (373), M45 (353), M46 (347), M47 (332) | G2 |
| | F36 (425) | M48 (338), M49 (364), M50 (370), M51 (390) | G3 |
| | F37 (354) | M48 (338), M49 (364), M50 (370), M51 (390) | G3 |
| | F38 (415) | M48 (338), M49 (364), M50 (370), M51 (390) | G3 |
| **Constance** | F12 (383) | M16 (317), M17 (266), M18 (287), M19 (297) | C1 |
| | F13 (371) | M16 (317), M17 (266), M18 (287), M19 (297) | C1 |
| | F14 (322) | M16 (317), M17 (266), M18 (287), M19 (297) | C1 |
| | F15 (438) | M20 (324), M21 (305), M22 (308), M23 (400) | C2 |
| | F16 (480) | M20 (324), M21 (305), M22 (308), M23 (400) | C2 |
| | F17 (334) | M20 (324), M21 (305), M22 (308), M23 (400) | C2 |
| | F18 (412) | M24 (340), M25 (425), M26 (383), M27 (382) | C3 |
| | F19 (330) | M24 (340), M25 (425), M26 (383), M27 (382) | C3 |
| | F20 (415) | M24 (340), M25 (425), M26 (383), M27 (382) | C3 |
| ***Introduced populations*** | | | |
| **Allos** | F1 (183) | M1 (183), M2 (201), M3 (229) | A1 |
| | F2 (194) | M1 (183), M2 (201), M3 (229) | A1 |
| | F3 (214) | M4 (194), M5 (193), M6 (229) | A2 |
| | F4 (174) | M4 (194), M5 (193), M6 (229) | A2 |
| | F5 (204) | M7 (285), M8 (292), M9 (266) | A3 |
| | F6 (220) | M7 (285), M8 (292), M9 (266) | A3 |
| | F7 (213) | M10 (212), M11 (224), M12 (195) | A4 |
| | F8 (195) | M10 (212), M11 (224), M12 (195) | A4 |
| **Pavin** | F21 (266) | M28 (272), M29 (240), M30 (288), M31 (255) | P1 |
| | F22 (262) | M28 (272), M29 (240), M30 (288), M31 (255) | P1 |
| | F23 (267) | M28 (272), M29 (240), M30 (288), M31 (255) | P1 |
| | F25 (288) | M32 (273), M33 (237), M34 (245), M35 (276) | P2 |
| | F26 (280) | M32 (273), M33 (237), M34 (245), M35 (276) | P2 |
| | F27 (246) | M36 (296), M37 (261), M38 (248), M39 (257) | P3 |
| | F28 (268) | M36 (296), M37 (261), M38 (248), M39 (257) | P3 |
| | F29 (265) | M36 (296), M37 (261), M38 (248), M39 (257) | P3 |

TL refers to total length (mm).

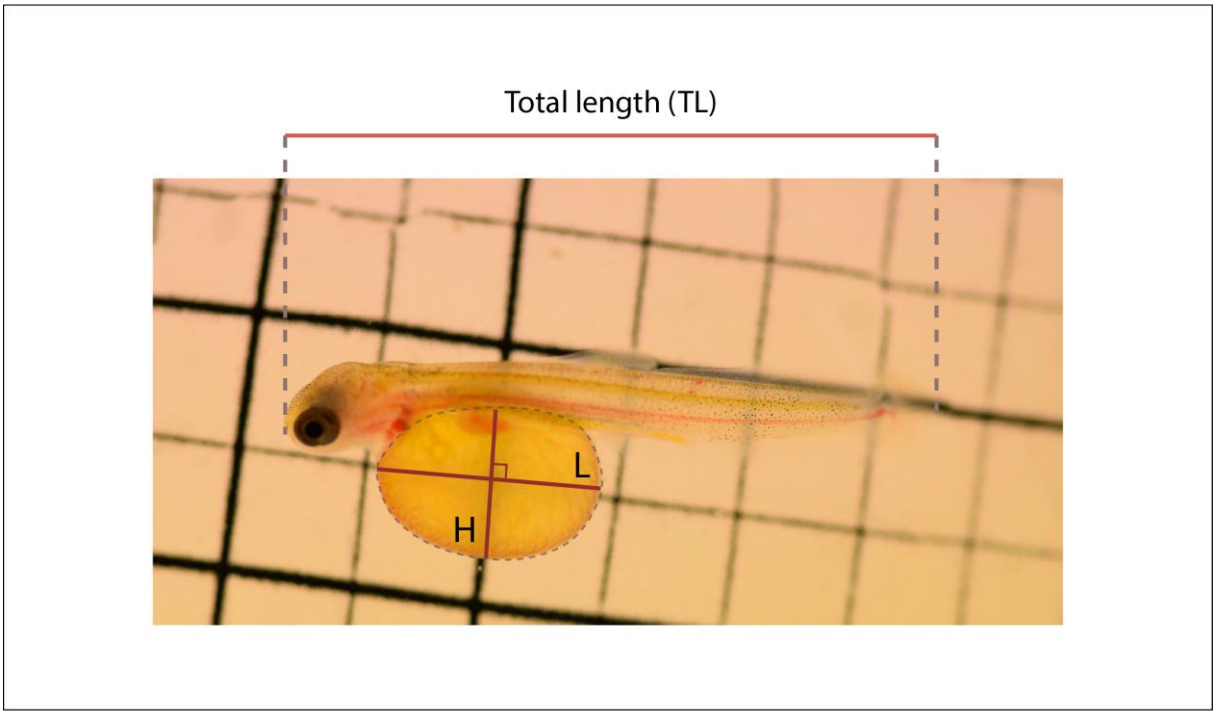


**Figure S1.** Morphometric measurements of Arctic charr (*Salvelinus alpinus*) larvae at hatching used to estimate yolk sac volume (YSV). Total length (TL) was measured from the tip of the snout to the end of the tail. Yolk sac length (L) and yolk sac height (H) correspond to the longest and shortest axes of the yolk sac, respectively.

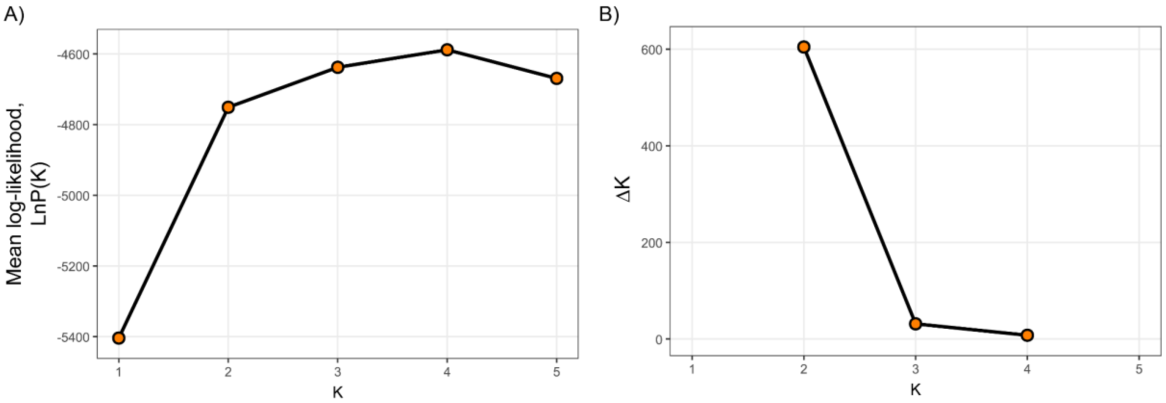


**Figure S2.** Determination of the number of genetic clusters and population genetic structure in Arctic charr (*Salvelinus alpinus*). A, mean log-likelihood values (LnP(K)) and B, ΔK values calculated as a function of the number of genetic clusters (K), averaged over 10 independent STRUCTURE runs per K. The Evanno method indicates a clear maximum at K = 2.

**Table S3.** Neutral genetic diversity at seven microsatellite loci across four Arctic charr (*Salvelinus alpinus*) populations.

| Population | | Cocl3_N | OtsG253b | Ssa456 | Sfo8 | *Sfo23* | Ssa85 | Omy301 | Overall |
|---|---|---|---|---|---|---|---|---|---|
| ***Native populations*** | | | | | | | | | |
| ***Geneva*** | ***N*** | 20 | 20 | 20 | 20 | 20 | 20 | 20 | 20 |
| | ***A*** | 8 | 6 | 4 | 8 | 7 | 12 | 7 | 52 |
| | ***Ar*** | 8 | 6 | 4 | 8 | 7 | 12 | 7 | 7.43 |
| | ***Ho*** | 0.65 | 0.55 | 0.65 | 0.55 | 0.35 | 0.8 | 0.6 | 0.59 |
| | ***He*** | 0.78 | 0.69 | 0.68 | 0.73 | 0.69 | 0.78 | 0.79 | 0.73 |
| | ***Fis*** | 0.16 | 0.2 | 0.05 | 0.25 | 0.49 | -0.02 | 0.24 | 0.20 [0.09-0.31] |
| | ***HWE*** | 0.00 | 0.39 | 0.39 | 0.00 | 0.00 | 0.44 | 0.01 | 0.57 |
| ***Constance*** | ***N*** | 20 | 20 | 20 | 20 | 20 | 20 | 20 | 20 |
| | ***A*** | 14 | 8 | 12 | 11 | 11 | 18 | 9 | 83 |
| | ***Ar*** | 14 | 8 | 12 | 11 | 11 | 18 | 9 | 11.86 |
| | ***Ho*** | 0.75 | 0.85 | 0.9 | 0.45 | 0.65 | 0.85 | 0.45 | 0.7 |
| | ***He*** | 0.92 | 0.84 | 0.9 | 0.89 | 0.8 | 0.92 | 0.84 | 0.87 |
| | ***Fis*** | 0.19 | -0.01 | 0 | 0.49 | 0.18 | 0.08 | 0.46 | 0.20 [0.06-0.35] |
| | ***HWE*** | 0.19 | 0.35 | 0.64 | 0.00 | 0.05 | 0.19 | 0.00 | 0.29 |
| ***Introduced populations*** | | | | | | | | | |
| ***Allos*** | ***N*** | 111 | 111 | 111 | 111 | 111 | 111 | 111 | 111 |
| | ***A*** | 13 | 9 | 6 | 15 | 10 | 14 | 10 | 77 |
| | ***Ar*** | 9.19 | 6.31 | 5.64 | 8.93 | 7.6 | 9.88 | 6.72 | 7.75 |
| | ***Ho*** | 0.46 | 0.83 | 0.67 | 0.67 | 0.41 | 0.77 | 0.59 | 0.63 |
| | ***He*** | 0.8 | 0.74 | 0.69 | 0.75 | 0.74 | 0.81 | 0.65 | 0.74 |
| | ***Fis*** | 0.43 | -0.12 | 0.04 | 0.11 | 0.44 | 0.05 | 0.09 | 0.15 [0.02-0.29] |
| | ***HWE*** | 0.00 | 0.04 | 0.07 | 0.00 | 0.00 | 0.41 | 0.01 | 0.57 |
| ***Pavin*** | ***N*** | 25 | 25 | 25 | 25 | 25 | 25 | 25 | 25 |
| | ***A*** | 13 | 9 | 11 | 7 | 12 | 14 | 6 | 72 |
| | ***Ar*** | 12.31 | 8.76 | 10.7 | 6.79 | 10.93 | 12.75 | 5.96 | 9.74 |
| | ***Ho*** | 0.64 | 0.92 | 0.92 | 0.8 | 0.48 | 0.88 | 0.48 | 0.73 |
| | ***He*** | 0.9 | 0.84 | 0.86 | 0.77 | 0.84 | 0.9 | 0.79 | 0.84 |
| | ***Fis*** | 0.29 | -0.09 | -0.07 | -0.04 | 0.43 | 0.02 | 0.39 | 0.13 [0.00-0.29] |
| | ***HWE*** | 0.00 | 0.08 | 0.07 | 0.06 | 0.00 | 0.81 | 0.00 | 0.43 |

N, number of genotyped individuals, A, number of alleles, Ar, allelic richness rarefied to the minimum sample size across populations, Ho, observed heterozygosity, He, expected heterozygosity, FIS, inbreeding coefficient estimated following Weir & Cockerham, HWE, Hardy-Weinberg equilibrium exact test p-values for each locus with the overall value corresponding to the proportion of loci deviating from HWE after Benjamini–Hochberg correction. The Overall column summarizes population-level estimates with N representing the mean number of genotyped individuals per locus, A the total number of alleles across loci, and Ar, Ho, He, and FIS the mean values across loci. For FIS, 95% confidence intervals were estimated by bootstrapping loci (10,000 iterations). Because Ar was rarefied to the minimum sample size across populations (N = 20), Ar equals A for Geneva and Constance. Ar estimates were robust to sample size imbalance, as confirmed by 10,000 balanced subsamplings. Mean Ar ± SD from subsampling was 7.54 ± 0.40 for Allos and 9.71 ± 0.28 for Pavin.

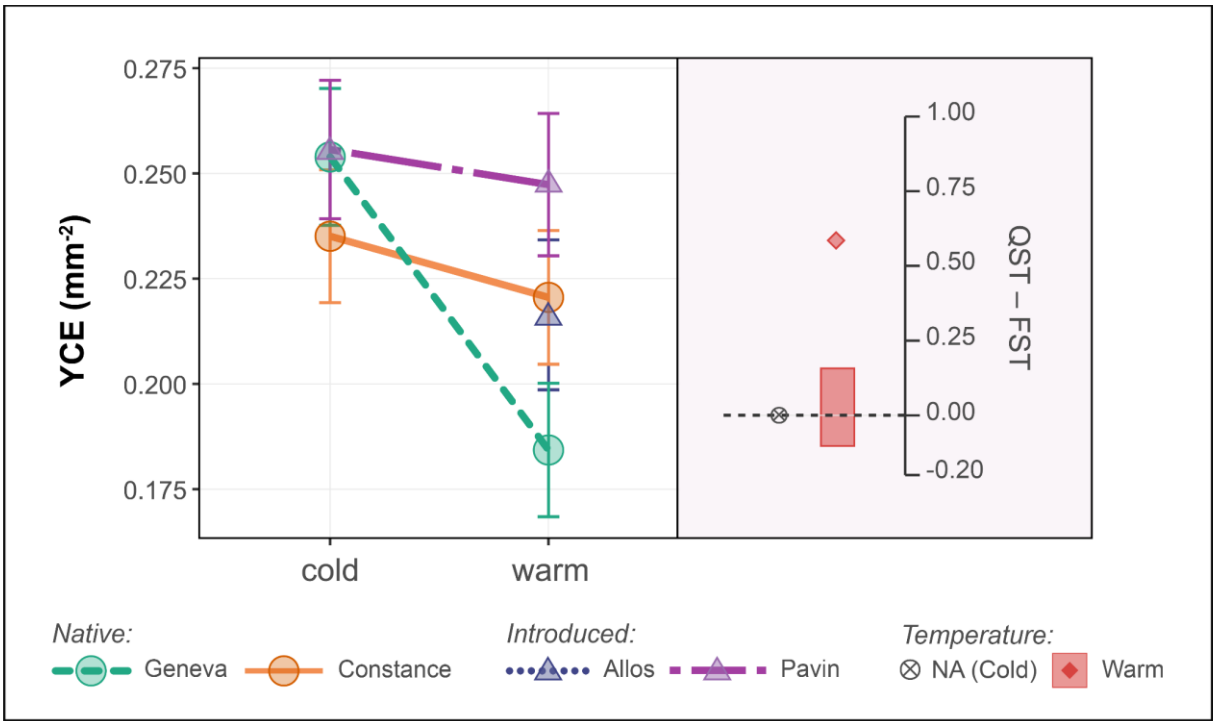


**Figure S3.** Thermal reaction norms and $Q_{st} - F_{st}$ estimates for yolk sac conversion efficiency (YCE) in four Arctic charr populations reared under cold (5 °C) and warm (8.5 °C) conditions. Left panel shows model-estimated marginal means (± 95% CI) of YCE (mm/mm$^3$). Right panel displays corresponding $Q_{st} - F_{st}$ values. Rectangles indicate the simulated neutral expectation. Filled symbols denote significant deviations from neutrality ($p < 0.05$). $Q_{st}$ could not be estimated under cold conditions due to the absence of additive genetic variance. Circles indicate native warm-origin populations (Geneva: green, Constance: orange), whereas triangles indicate introduced cold-origin populations (Allos: blue, Pavin: purple).

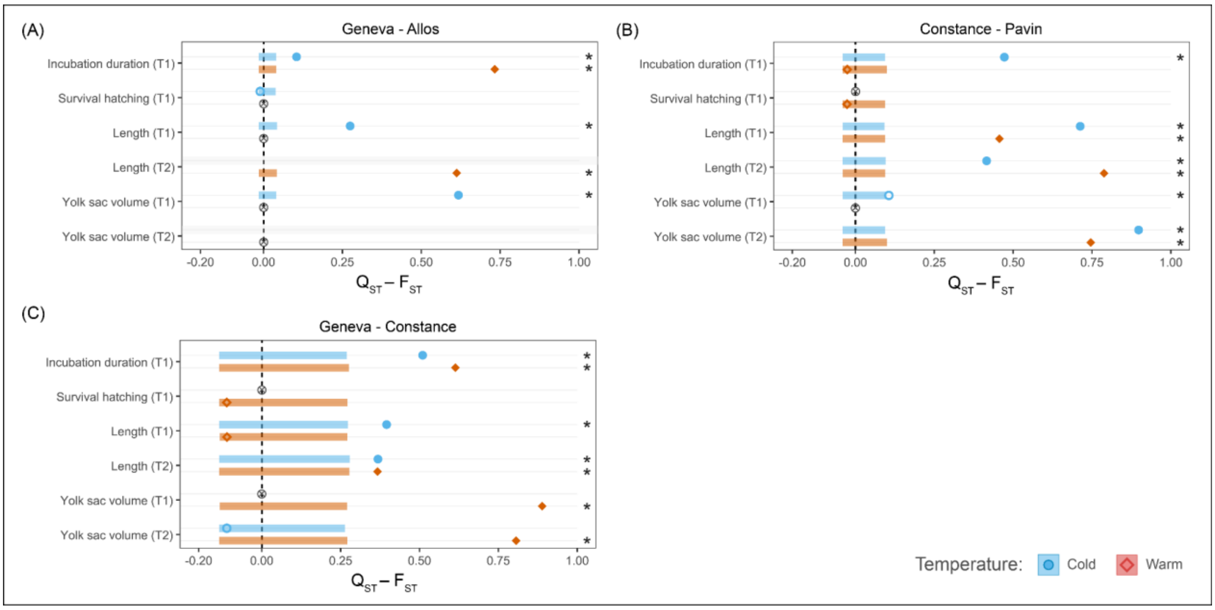


**Figure S4.** Simulated neutral expectations and observed values of $Q_{st} - F_{st}$ for population comparisons in Arctic charr (*Salvelinus alpinus*). A, Geneva–Allos, B, Constance–Pavin, and C, Geneva–Constance. Observed values represent $Q_{st} - F_{st}$ for incubation duration, survival, body length and yolk sac volume measured at hatching (T1) and at the end of the experiment (T2), under cold (blue circle) and warm (orange diamonds) rearing conditions. Rectangles represent the neutral expectation interval of $Q_{st} - F_{st}$, simulated following Whitlock and Guillaume (2009), based on empirical estimates of $F_{st}$ and within-population variance for each trait. Filled symbols indicate significant deviations from neutrality ($p < 0.05$), whereas open symbols indicate non-significant deviations. Crossed circles correspond to traits for which the test was not applicable due to the absence of between-population variance or within-population variance.

**Table S4.** Summary of the linear mixed models and generalized linear mixed-effects models assessing the variance components and $Q_{st}$ estimates for quantitative traits and early survival in *Salvelinus alpinus*.

| | *Cold* | | | | | *Warm* | | | | |
|---|---|---|---|---|---|---|---|---|---|---|
| | $\sigma_{GB}{}^2$ | $\sigma_{GW}{}^2$ | $Q_{st}$ | *p-val.* | *h²* | $\sigma_{GB}{}^2$ | $\sigma_{GW}{}^2$ | $Q_{st}$ | *p-val.* | *h²* |
| ***Overall*** | | | | | | | | | | |
| Incubation duration | 507.93 | 96.13 | 0.73 | < 0.001 | 0.88 [0.78-0.92] | 790.49 | 118.8 | 0.77 | < 0.001 | 0.77 [0.59-0.86] |
| Survival at hatching | 0.02 | 0.04 | 0.21 | 0.07 | 0.00 [0.00-0.06] | 0.14 | 0.29 | 0.20 | 0.09 | 0.07 [0.00-0.14] |
| Length (T1) | 0.17 | 0.06 | 0.59 | < 0.001 | 0.26 [0.12-0.39] | 0.16 | 0.02 | 0.79 | < 0.001 | 0.08 [0.00-0.30] |
| Length (T2) | 0.09 | 0.17 | 0.21 | 0.10 | 0.28 [0.08-0.43] | 1.50 | 0.32 | 0.70 | < 0.001 | 0.21 [0.04-0.35] |
| YSV (T1) | 5.17 | 2.67 | 0.49 | < 0.001 | 0.07 [0.00-0.21] | 3E-08 | 2.46 | 0.00 | 0.98 | 0.04 [0.00-0.24] |
| YSV (T2) | 3.23 | 0.85 | 0.66 | < 0.001 | 0.12 [0.00-0.25] | 25.3 | 2.72 | 0.82 | < 0.001 | 0.11 [0.00-0.23] |
| YCE | 2E-04 | 0.00 | – | – | – | 5E-04 | 1E-04 | 0.69 | < 0.001 | 0.12 [0.00-0.60] |
| ***Geneva - Allos*** | | | | | | | | | | |
| Incubation duration | 8.52 | 32.38 | 0.12 | < 0.01 | – | 1925.39 | 162.4 | 0.86 | < 0.001 | – |
| Survival at hatching | 0.000 | 18.27 | – | – | – | 0.09 | 0.00 | – | – | – |
| Length (T1) | 0.027 | 0.03 | 0.29 | < 0.001 | – | 0.00 | 0.00 | – | – | – |
| Length (T2) | | | | | – | 0.66 | 0.20 | 0.62 | < 0.001 | – |
| YSV (T1) | 12.86 | 3.80 | 0.63 | < 0.001 | – | – | – | – | – | – |
| YSV (T2) | | | | | – | 0.54 | 0.00 | – | – | – |
| ***Constance - Pavin*** | | | | | | | | | | |
| Incubation duration | 307.31 | 154.4 | 0.50 | < 0.001 | – | 0.00 | 190.6 | – | 0.85 | – |
| Survival at hatching | 1E-04 | 4E-09 | - | - | – | 0.00 | 0.43 | – | – | – |
| Length (T1) | 0.42 | 0.07 | 0.74 | < 0.001 | – | 0.16 | 0.08 | 0.48 | < 0.001 | – |
| Length (T2) | 0.29 | 0.18 | 0.44 | < 0.001 | – | 3.79 | 0.43 | 0.81 | < 0.001 | – |
| YSV (T1) | 0.51 | 1.67 | 0.13 | < 0.001 | – | 0.00 | 6.52 | 0.00 | 0.84 | – |
| YSV (T2) | 19.793 | 0.81 | 0.92 | < 0.001 | – | 51.07 | 7.52 | 0.77 | < 0.001 | – |
| ***Geneva - Constance*** | | | | | | | | | | |
| Incubation duration | 179.8 | 54.8 | 0.62 | < 0.001 | – | 866 | 165.2 | 0.72 | < 0.001 | – |
| Survival at hatching | 0.00 | 3E-12 | – | – | – | 0.000 | 0.15 | – | – | – |
| Length (T1) | 0.09 | 0.04 | 0.51 | < 0.001 | – | 3E-07 | 0.02 | 0.00 | 0.88 | – |
| Length (T2) | 0.39 | 0.21 | 0.48 | 0.01 | – | 0.938 | 0.51 | 0.48 | 0.01 | – |
| YSV (T1) | 0.00 | 2.22 | 0.00 | 0.88 | – | 7.384 | 0.00 | 1.00 | < 0.001 | – |
| YSV (T2) | 2E-04 | 1.03 | 0.00 | 0.88 | – | 59.37 | 2.68 | 0.92 | < 0.001 | – |

Between-population additive genetic variance ($\sigma_{GB}{}^2$), within-population additive genetic variance estimated as four times the sire variance ($\sigma_{GW}{}^2 = 4 \times \sigma_{sire}{}^2$) are reported. Narrow-sense heritability ($h^2$) was estimated overall as the ratio of additive genetic variance to total phenotypic variance. Values are reported with their 95% confidence intervals obtained by parametric bootstrap (1,000 simulations). Significant $Q_{st}$ estimates are indicated in bold. YSV = yolk sac volume, YCE = yolk sac conversion efficiency.
